# Using accelerometers to infer behaviour of cryptic species in the wild

**DOI:** 10.1101/2023.03.20.533342

**Authors:** Laura Benoit, Nadège C. Bonnot, Lucie Debeffe, David Grémillet, A.J. Mark Hewison, Pascal Marchand, Laura Puch, Arnaud Bonnet, Bruno Cargnelutti, Nicolas Cebe, Bruno Lourtet, Aurélie Coulon, Nicolas Morellet

## Abstract

Accelerometery is revolutionising the field of behavioural ecology through its capacity to detect the fine-scale movements of animals resulting from their behaviour. Because it is often difficult to infer the behaviour of wildlife on a continuous basis, particularly for cryptic species, accelerometers potentially provide powerful tools for remote monitoring of their behavioural responses to the environment.

The goal of this study was to provide a detailed, calibrated methodology, including practical guidelines, to infer the behaviour of free-ranging animals from acceleration data. This approach can be employed to reliably infer the time budget of species that are difficult to observe in certain environments or at certain times of the day. To this end, we trained several behavioural classification algorithms with accelerometer data obtained on captive roe deer, then validated these algorithms with data obtained on free-ranging roe deer, and finally predicted the time-budgets of a substantial sample of unobserved free-ranging roe deer in a human-dominated landscape.

The best classification algorithm was the Random Forest which predicted five behavioural classes with a high overall level of accuracy (≈ 90%). Except for grooming (34-38%), we were able to predict the behaviour of free-ranging roe deer over the course of a day with high accuracy, in particular, foraging head down, running, walking and immobile (68-94%). Applied to free-ranging individuals, the classification allowed us to estimate, for example, that roe deer spent about twice as much time foraging head-down, walking or running during dawn and dusk than during daylight or night-time.

By integrating step by step calibration and validation of accelerometer data prior to application in the wild, our approach is transferable to other free-ranging animals for predicting key behaviours in cryptic species.

## Introduction

Behaviour is increasingly recognised as a fundamental component of life history (Wolf *et al*. 2007) and a key mechanism that enables wild populations to cope with global change (Caro 1999). However, obtaining behavioural data for a large number of individuals in the wild is a major challenge, particularly for elusive species. Direct observation is often constrained by visibility (Löttker *et al*. 2009; Brown *et al*. 2013), which may vary over the day (e.g. lower during night-time), with meteorological conditions (e.g. lower during rainy/foggy days), among habitats (e.g. forest vs open habitat), or individuals (e.g. lower for shy individuals). Moreover, the presence of an observer may modify the behaviour of the focal animal and may thus bias studies relying on direct observation (Schneirla 1950; Tuyttens *et al*. 2014). The development of positioning devices (Kooyman 2004) such as VHF, Argos or GPS-tracking has revolutionized the field of behavioural ecology (Cagnacci *et al*. 2010; Gurarie *et al*. 2016), providing detailed information on the movement (e.g. speed or velocity: Ponganis *et al*. 1990; Malagnino *et al*. 2021) and spatial behaviour (e.g. daily space use: Seigle-Ferrand *et al*. 2020, migration: Dujon *et al*. 2017; dispersal: Cozzi *et al*. 2020) of wild animals. However, these devices provide incomplete information, in the sense that we know *where the animal is* but not *what it is doing*. More recently, with further technological progress, animal-borne sensors (Cooke *et al*. 2004; Whitford & Klimley 2019) can now collect information on body movements (accelerometer: Yoda *et al*. 1999; Brown *et al*. 2013), internal temperature (stomach temperature biologgers: Wilson *et al*. 1995; Weimerskirch *et al*. 2005), heart rate (cardiac biologger: Grémillet *et al*. 2005; Ditmer *et al*. 2018) and stress hormones (blood sampler: Ponganis *et al*. 1997; Takei *et al*. 2016).

Accelerometer biologgers are promising and powerful tools to access detailed information on behaviour. Acceleration is measured along two or three axes, with sampling frequencies usually varying from 8 to 100 hertz (i.e. 8 to 100 measurements per second: Brown *et al*. 2013), providing detailed and continuous information on fine-scale body movements and posture (Sato 2003; Shepard *et al*. 2008b). First developed for domestic and aquatic animals (Yoda *et al*. 1999; Watanabe *et al*. 2005), accelerometers have since been adapted for wild terrestrial species (Brown *et al*. 2013), generating an incredible amount of information regarding animal state (Wilson *et al*. 2014), behaviour (Graf *et al*. 2015), associated energy expenditure (Wilson *et al*. 2006; Mosser *et al*. 2014), and fitness components (Grémillet *et al*. 2018; Marchand *et al*. 2021a).

To accurately infer behaviour from accelerometer data, supervised machine learning methods are frequently employed (Sakamoto *et al*. 2009; Nathan *et al*. 2012; Tatler *et al*. 2018). These approaches use classification algorithms to learn the relationship between the acceleration data and observed behaviours (calibration), so that when algorithm performance is high (validation), it can successfully predict behaviour with novel acceleration data (application). A key requirement is, thus, a behaviour labelled-accelerometer dataset where the labels are derived from films or direct observations of wild animals (Kröschel *et al*. 2017), animals equipped with video cameras (Volpov *et al*. 2015), captive animals (Graf *et al*. 2015; Rast *et al*. 2020) or surrogate species (Pagano *et al*. 2017; Ferdinandy *et al*. 2020).

These methods have been successfully applied to a variety of species (Brown *et al*. 2013) to identify routine behaviours such as walking, flying, or swimming (Shepard *et al*. 2008b), but also to detect specific events such as prey capture (Ropert-Coudert *et al*. 2006; Wilson *et al*. 2013), mating or suckling (Whitney *et al*. 2010; Shuert *et al*. 2018). Thus, it is possible to infer behaviour when visibility is limited, providing a detailed insight into daily time budgets (Lush *et al*. 2016; Rast *et al*. 2020), even for elusive species.

The aim of our study is to present practical guidelines for inferring animal behaviour in the wild, based on a classification algorithm for acceleration data trained on captive animals, to calibrate the approach for species that are particularly elusive. For this purpose, we studied a large herbivore, the roe deer (*Capreolus capreolus*) using both captive individuals in an enclosure and free-ranging individuals monitored in a natural environment. Indeed, the roe deer is typically elusive, and predominantly crepuscular or nocturnal, particularly when hunted (Bonnot et al. 2020), so that direct observation is both time consuming and often incomplete, particularly in closed habitats. Although its spatial behaviour and activity have been intensively studied independently (Bonnot et al. 2013; Padié et al. 2015; Krop-Benesch et al. 2013), to provide a more complete and unfragmented picture of both where the animal is and what it is doing there, we need to be able to link spatial locations (e.g. GPS monitoring) with continuous inference of behaviour. Accelerometers provide an unprecedented opportunity to infer behavioural states over the 24-hour cycle to understand, for instance, how cryptic species behave across fluctuating environments.

To this end, i) we calibrated a behavioural classification algorithm using supervised machine learning methods (see Material & Methods), accelerometer data derived from sensors integrated in GPS-collars, and behavioural videos of captive roe deer (Calibration step, Fig.1), ii) we validated algorithm predictions on free-ranging roe deer with the same type of accelerometers and filmed in the wild (Validation step) and finally iii) to illustrate the relevance of this approach, we applied this algorithm to predict the behaviours of 47 unobserved free-ranging roe deer with accelerometers to infer their circadian time budget in a human-dominated landscape (Application step).

**Figure 1:**
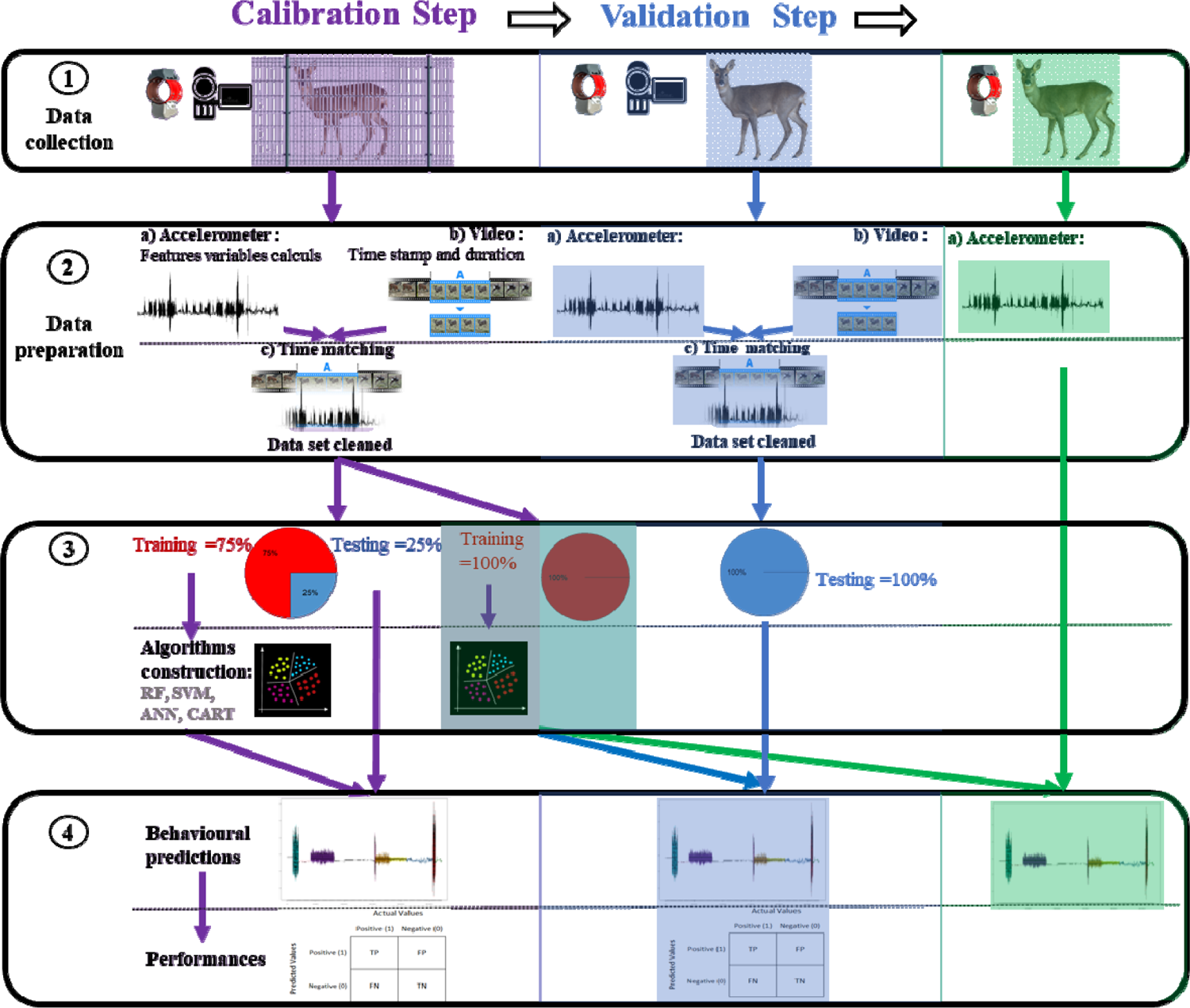
Flowchart of the three-step process developed for the classification of roe deer behaviour using accelerometer data. Calibration step (on filmed captive roe deer), Validation step (on filmed free-ranging roe deer), Application step (on unobserved free-ranging roe deer). The four components of analysis (numbered 1 to 4) were described for the calibration step column and then mentioned in the two other columns if they were still running: Part IZ: Collection of behavioural and accelerometer data on roe deer. Part IZ: Preparation of behavioural (time stamp and duration of behavioural sequences) and accelerometer (calculation of 26 feature variables) data, and temporal synchronisation between the two. Part IZ: Construction of four classification algorithms based on the training data subset. Part IZ: Prediction of behaviours from classification algorithms and estimation of performance based on the test data subset.

## Materials and Methods

### Accelerometer characteristics

Roe deer were equipped with one of two types of accelerometry sensors associated with two different models of GPS collars (see Table 1). Both sensor types measured acceleration along the three perpendicular axes (anteroposterior [surge or forward/backward], dorso-ventral [heave or up/down] and transversal [sway or sideways], hereafter called X, Y and Z axes, respectively) with values ranging between −125 (= −8 g) and 125 (= 8 g) for each axis (1 g = 9.81 m.s^-2^). However, although they were supplied by the same manufacturer, the two sensor types differed in their acceleration pattern due to i) a difference in sampling frequency (i.e. 8 Hz and 32 Hz), ii) their position on the animal’s neck (due to weight differences) and iii) the orientation of the axes on the collar (see images in Table 1). To account for these differences, we constructed distinct classification algorithms for the two accelerometer sensor types (type A or B). In addition, to control for inter-individual variation in acceleration patterns for a given type of sensor (related to collar position on the animal’s neck or variation in sensitivity between sensors; Kröschel *et al*. 2017; Dickinson *et al*. 2020), we scaled acceleration data at each step (see details in section 2.b).

**Table 1:**
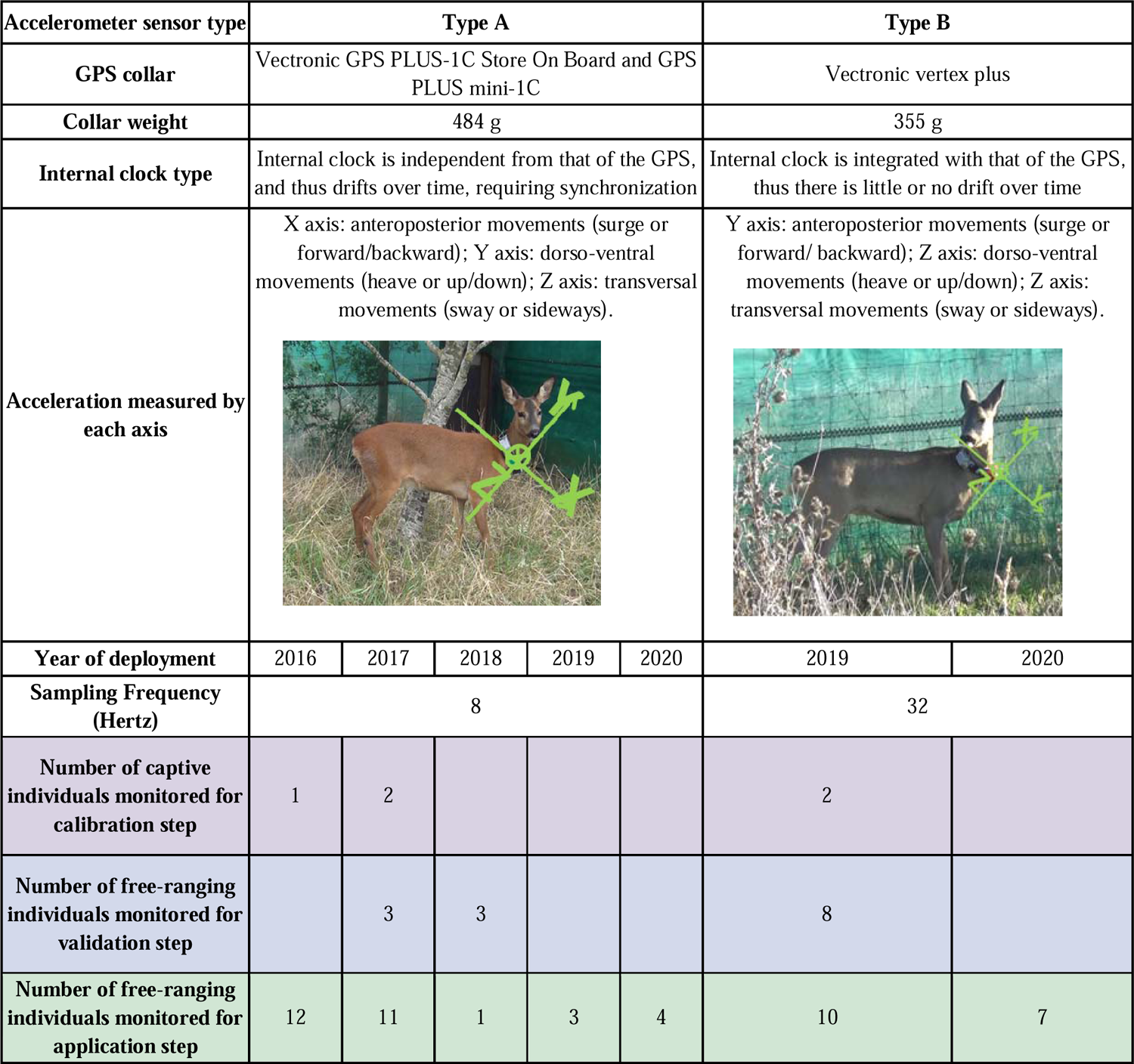
Details of sensor characteristics and number of individuals monitored at each step (Calibration, Validation, Application) of the study

| Accelerometer sensor type | Type A |  |  |  |  | Type B |  |
| --- | --- | --- | --- | --- | --- | --- | --- |
| GPS collar | Vectronic GPS PLUS-1C Store On Board and GPS PLUS mini-1C |  |  |  |  | Vectronic vertex plus |  |
| Collar weight | 484 g |  |  |  |  | 355 g |  |
| Internal clock type | Internal clock is independent from that of the GPS, and thus drifts over time, requiring synchronization |  |  |  |  | Internal clock is integrated with that of the GPS, thus there is little or no drift over time |  |
| Acceleration measured by each axis | X axis: anteroposterior movements (surge or forward/backward); Y axis: dorso-ventral movements (heave or up/down); Z axis: transversal movements (sway or sideways). |  |  |  |  | Y axis: anteroposterior movements (surge or forward/ backward); Z axis: dorso-ventral movements (heave or up/down); Z axis: transversal movements (sway or sideways). |  |
| Year of deployment | 2016 | 2017 | 2018 | 2019 | 2020 | 2019 | 2020 |
| Sampling Frequency (Hertz) | 8 |  |  |  |  | 32 |  |
| Number of captive individuals monitored for calibration step | 1 | 2 |  |  |  | 2 |  |
| Number of free-ranging individuals monitored for validation step |  | 3 | 3 |  |  | 8 |  |
| Number of free-ranging individuals monitored for application step | 12 | 11 | 1 | 3 | 4 | 10 | 7 |

#### I) Calibration of the classification algorithms on captive roe deer

In this first step, we built and evaluated the performance of behavioural classification algorithms trained on accelerometer data and behavioural observations (Fig.1). For that, we studied captive roe deer that were easily observable in the experimental facility of INRAE in order to obtain numerous high-quality video data (Fig.1).

### 1) Collection of behavioural and accelerometer data

The Gardouch experimental station is located in the South-West of France (43°37′N, 1°67′E), on the slopes of a hill, experiencing an oceanic climate with summer droughts. The station includes 11 enclosures of 0.5 ha which house tame captive roe deer of both sexes (one to six roe deer per enclosure). Each enclosure contains grassland, trees and shrubs that provide shelter and some natural food sources, but their diet is supplemented with artificial pellets for livestock (600 g per individual per day).

Accelerometer data were collected between 2016 and 2019 on three tame adult females living in these enclosures with one or three congeners. Because they are accustomed to human presence, these deer can be fitted with collars without requiring physical restraint and, subsequently, easily observed. This protocol was approved by the Ethical Committee 115 of Toulouse and was authorized by the French government (APAFIS#15760-2018061909204934 v6). In order to maximise the range of observed behaviours, the accelerometers were deployed on each individual several times, for periods of approximately 2-weeks, during different seasons and years. Furthermore, two females were fitted with both sensor types across years (see Table S2). Behavioural observations (see below) were collected by video-recording each female during daytime, while they moved freely within their enclosure. These recordings were time-stamped manually (GMT-time) using a GPS-beacon to provide exact synchronisation with the accelerometer data.

### 2) Data preparation

#### 2.a) Behavioural observations from video recording

A very detailed ethogram of 54 exclusive behaviours was defined, based on biologically meaningful behaviours that involve specific positions of the body and head (see Appendix S1). When an animal could not be observed, or when the distinction between behaviours was unclear, we classified the event as “unwatchable” and removed these sequences from the dataset. In order to obtain the time stamp and duration of each behaviour, video analyses were carried out using freely available open-source event-logging software (“Jwatcher”and “BORIS”: Stankowich 2008; Friard & Gamba 2016). Each behavioural event was manually labelled using the pre-defined ethogram. Because certain behaviours were not expressed sufficiently frequently by the captive animals (ex: fighting), we did not consider them for the subsequent classification step (more information on behavioural data obtained per individual are provided in Appendix S2). Moreover, we grouped certain behaviours that are very similar in terms of movement and function (e.g. observing and sniffing the air) into a single category, resulting in 25 behavioural classes for the classification algorithms (see Tables S1 and S2 for more details).

#### 2.b) Accelerometer data

We derived 26 separate feature variables (see Table 2 and Collins *et al*. (2015), for details) from acceleration signals measured on the three axes (Table 1). In particular, for each axis, we calculated standard deviation, static acceleration, dynamic acceleration, angle of rotation, minimum and maximum values for dynamic acceleration, dominant power spectrum and frequency at the dominant power spectrum. In addition, based on all three axes, we calculated overall and vectorial body dynamic acceleration (OBDA and VEDBA), which provide proxies of energy expenditure linked to body movement (Qasem *et al*. 2012).

**Table 2:**
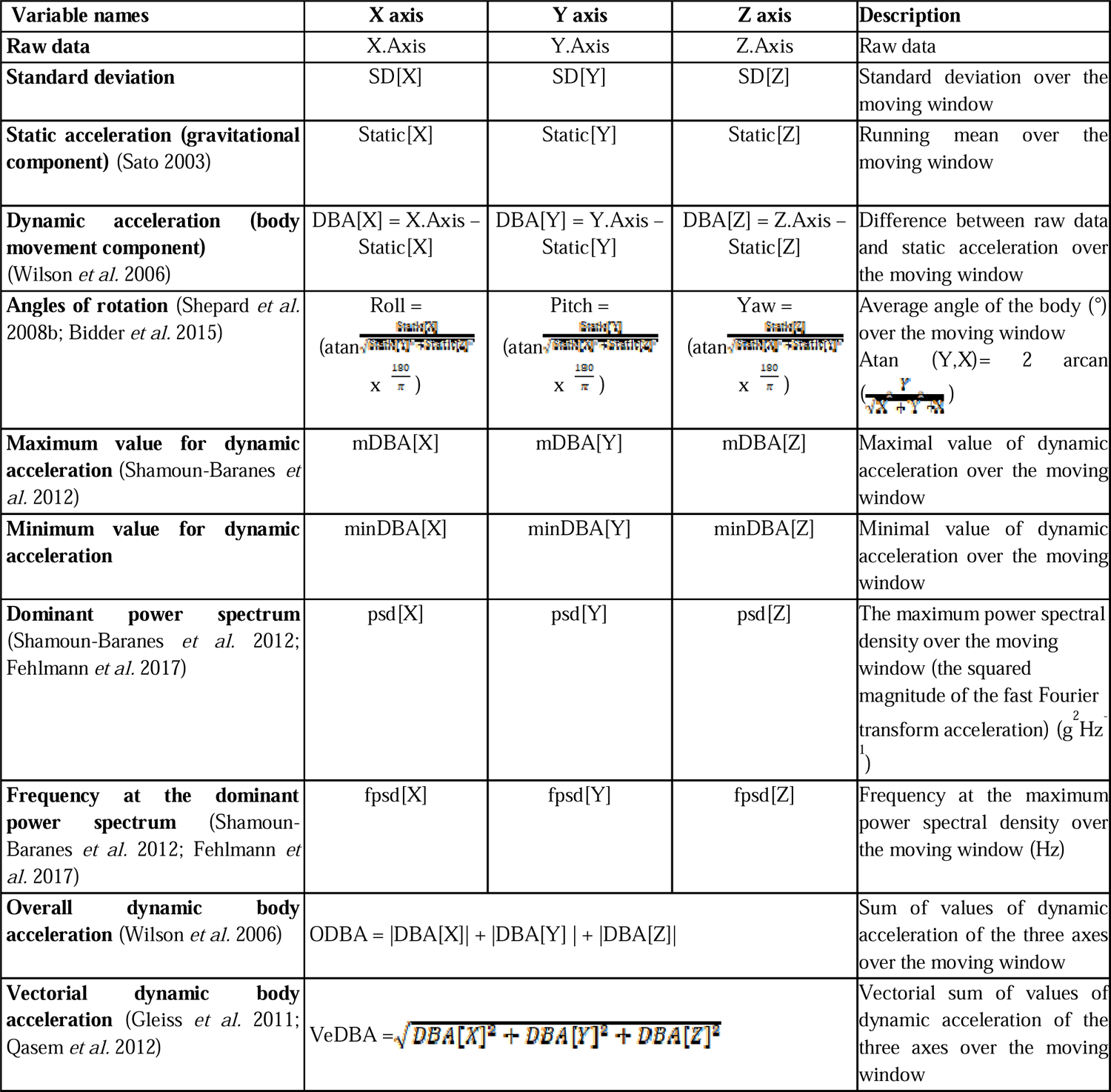
Details of the 26 variables derived from raw accelerometer data and used to infer roe deer behaviour. The duration of the moving window over which the variables were calculated was 4 seconds for the type A sensor and 3 seconds for the type B sensor (see main text for details).

The duration of the moving window over which these 26 variables were calculated was based on the accuracy of the classification algorithm independently for the two types of sensors. We tested moving windows of between 1 and 5 seconds (following Shepard *et al*. (2008a)), and obtained the best performance with a moving window of 4 seconds for the type A sensor and 3 seconds for the type B sensor, likely due to different base sampling frequencies (8 versus 32 Hz, respectively).

Subsequently, we scaled the acceleration data by subtracting the mean of the feature variable and dividing the result by its standard deviation. More specifically, for this calibration step and for each sensor type, we scaled data using the mean and standard deviation calculated on data from all captive individuals. Indeed, there was little among-individual variation in acceleration patterns across captive animals, probably because of small sample sizes (N=3 for type A and N=2 for type B), and/or because they were equipped for short time periods (one to two weeks) so that there was little variation in fur thickness.

#### 2.c) Synchronising acceleration data with observational data

An accurate calibration of the accelerometer signal requires a perfect match of the accelerometer data with the corresponding behavioural sequence derived from the video observations. However, initial inspection revealed that the internal clocks of the accelerometers often did not run at precisely the same rate as the GPS-beacon clock used during video-recording. We thus corrected, when possible, the time of the accelerometer data (see Appendix S2) and retained only synchronised data for further analyses.

Finally, we cleaned the dataset by retaining only behaviourally labelled accelerometer data that corresponded to the sequences of the 25 behaviours defined above. To obtain a clear signal for these behaviours, we retained only those segments that lasted more than 2 seconds, and then removed the initial and final 0.5 second of each segment to eliminate transitions between successive behaviours.

### 3) Construction of classification algorithms

For the calibration step, we randomly selected 75% of the sequences for each behaviour (independent behavioural sequences) from the cleaned dataset of captive roe deer to train our classification algorithms. We compared several supervised machine-learning methods which are based on different rules to classify data (see Table 3). For each method, tuning steps and tuned parameters varied, as did the combination of feature variables used to optimize the classification (see Table 3). Based on the most commonly-used classification approaches, we tested random forests (hereafter, RF), artificial neural networks (ANN), support vector machine (SVM) and classification and regression trees (CART) (see Nathan *et al*. (2012) for more details).

**Table 3:** Supervised machine learning methods tested in this study.

| Supervised machine learning methods | R packages used and tuning | Rationale of the approach |
| --- | --- | --- |
| Random forest (RF) | *Caret* (method = rf → ntree (=500) and mtry (=15) optimal) required *Random forest* | This method is based on several iterations of classification trees using a random subset of the data for each tree (here n=500) and a random subspace of predictor variables (here n=9) for each tree branch. The final prediction is the result of all trees combined by a majority rule thus limiting overfitting and being able to cope with unbalanced datasets (Breiman 2001). |
| Artificial neural networks (ANN) | *Caret* (method = nnet → size (=5) and decay (=0) optimal) | This method distributes the data in automata (neurons). These units are responsible for combining their information together to determine the value of the discrimination variable. It is from the connection of these units to each other that the ANN capacity for discrimination emerges. |
| Support Vector Machine (SVM) | *e1071* | This method searches for an optimal linear classifier (hyperplane) that will separate the data into classes and maximize the distance between these classes (margins) |
| Classification And Regression Tree (CART) | *Caret* (method = rpart2→ maxdepth= 3 for sensor B and 4 for sensor A) required *rpart* | This method uses a set of hierarchical decision rules developed to classify data. It is a decision tree where each fork is split by values of a predictor variable. |

### 4) Predictions and performance of classification algorithms

Using each of the classification algorithms trained on 75% of the cleaned data set collected on captive animals (training dataset), we predicted behaviour on the remaining 25% (test dataset). We compared the resulting predicted behaviours from the different algorithms with the observed behaviours extracted from videos, based on confusion matrices. We calculated several metrics (see details in Appendix S3) to assess the performance of the algorithms for each behaviour separately (especially the F1-score metric) and the overall performance of the algorithms (especially the accuracy metric).

Finally, we focused on specific behaviours that we considered biologically meaningful to subsequently explore variation in the circadian time budget of free-ranging roe deer during the Application step. To do so, we grouped behaviours that were similar, or difficult to distinguish, into five behavioural classes: 1) foraging head down (including feeding, walking or unmoving with head-down), 2) walking head up (including walking with head-up and head at mid-height or sniffing the air while moving), 3) running, (corresponding to trotting, jumping or galloping), 4) grooming (including grooming while either standing or lying, and shaking the body) and 5) unmoving (including other behaviours such as lying down, standing still, vigilance or feeding with head-up) (see Appendix S1). The same performance metrics as described in Appendix S3 were then calculated for each of these five behavioural classes.

### II) Validation of the classification algorithms in the wild

In this second step, we validated the classification algorithms that were calibrated on captive roe deer for predicting behaviour of free-ranging individuals. To do so, we used independent accelerometer and observation data collected on free-ranging roe deer that were comparable to those used for the subsequent Application step in the wild (Fig.1).

#### 1) Collection of behavioural and accelerometer data

We studied a free-ranging roe deer population monitored in the Aurignac study site located in a 19,000 ha rural region in the Vallons et Coteaux de Gascogne (Zone Atelier PyGar) in the south–west of France (43°16′N, 0°53′E). The landscape is heterogeneous and hilly, with a mix of crops (32% of the total area), meadows (37%), hedgerows (4%) and woodland (19%) (see Morellet *et al*. (2011) for more details).

Accelerometer data were collected between 2016 and 2020 on free-ranging roe deer that were caught during winter (from January to March) using drive nets. Deer were tranquilised (with an intramuscular injection of acepromazine (calmivet 3cc; Montané *et al*. 2003) and transferred to a wooden retention box to reduce stress and risk of injury. Each individual was aged and sexed and then equipped with a GPS collar with integrated accelerometer sensors. All capture and marking procedures were done in accordance with local and European animal welfare laws (prefectural order from the Toulouse Administrative Authority to capture and monitor wild roe deer and agreement no. A31113001 approved by the Departmental Authority of Population Protection;). Marked individuals were video-recorded during the following year when conditions were suitable for the collection of behavioural information (Appendix S2). Videos were time-stamped manually (GMT) using a GPS-beacon (GPS Garmin).

#### 2) Data preparation

The preparation of behavioural (a: labelling behavioural sequences) and accelerometer (b: calculation) data and the time-synchronisation between them (c) were performed as explained in section I.2. However, in this validation step on free-ranging roe deer, we scaled all accelerometer variables per individual (to minimize variations due to collar tightness and sensor type) using accelerometer data recorded over approximatively one month, close in time to the moment when the individual was observed (to minimize variation due to seasons). The duration of one month was estimated to be sufficient to estimate deer activity while being relatively easily tractable for these accelerometer bigdata.

#### 3) Construction of classification algorithms

To validate the behavioural predictions of the classification algorithms on free-ranging roe deer, for each sensor type separately, we used the full cleaned dataset collected on captive animals as the training dataset to construct the classification algorithms (hereafter, referred to as the global classification algorithm). As for the calibration step, we tested different supervised machine learning methods (RF, ANN, CART and SVM).

#### 4) Predictions and performance of classification algorithms

Using the global classification algorithms, we predicted behaviour for the full cleaned dataset collected on free-ranging roe deer, and evaluated their performance by comparing the predicted and observed behavioural classes with the metrics presented in section I.4.

### III) Application of the classification algorithms to predict the circadian time budget of free ranging animals

In this last step, we applied the global classification algorithms (one for each sensor type) to free-ranging roe deer that were fitted with an accelerometer sensor, but that were not video recorded, in order to estimate their circadian time budget (Fig.1).

#### 1) Data collection, preparation and prediction of algorithms

We collected accelerometer data on 47 free-ranging roe deer from the Aurignac site as explained in section II.1. We focused this analysis on data collected during March, because there are no important life history events (e.g. birth, rut, etc.) at this time that might confound signal clarity, although males may begin defending territories. We calculated feature variables as described above (see section I.2), and scaled them per individual using all the data available during March for that individual. Then, we applied the global classification algorithms that we constructed on captive roe deer and validated on free-ranging roe deer (see section II.3) to obtain predictions per individual for each behavioural class for the month of March (see Fig.1).

#### 2) Estimation of circadian time budget

As the activity of roe deer is known to vary strongly over the circadian cycle, with higher activity during twilight (Krop-Benesch *et al*. 2013), we analysed the proportion of time allocated to each behavioural class both per day and per hour using Bayesian multilevel mixed models (with a Dirichlet distribution and a logit link function; using “brm” function in the brms package (Bürkner 2017)). First, we investigated the proportion of time allocated to each behavioural class per day with a generalized multilevel mixed model. Secondly, with a multilevel generalized additive mixed model, we investigated variation in the proportion of time allocated to each behavioural class per hour, in relation to the time of day, using cyclic cubic splines to smooth the effect of time (in hours) over the 24-hour cycle. We included individual identity as a random factor in all models to account for repeated observations for a given individual. Finally, we compared this model with a basic model with no effect of the time of day, based on approximate leave-one-out (LOO) cross-validation, as implemented in the loo package (Vehtari *et al*. 2017).

All analyses were performed with R version 4.0.3 (R Core Team, 2020).

## Results

### a) Comparing supervised machine learning approaches

For the calibration step, all supervised machine learning approaches predicted (i) the 25 behaviours that made up the test sub-dataset with an accuracy varying between 50% and 60% for type A sensors, and between 53% and 69% for type B, and (ii) the five behavioural classes with a high level of accuracy (from 85% to 91% for type A, and from 85% to 94% for type B) (see Appendices S3 & S5). For the validation step, all supervised machine learning methods predicted (i) the 25 behaviours that made up the test sub-dataset with an accuracy varying between 30% and 46% for type A sensors, and between 36% and 48% for type B, and (ii) the five behavioural classes with a high level of accuracy (from 79% to 90% for type A, and from 58% to 86% for type B) (see Appendices S8 & S10). Overall, when considering the ethogram of 25 behaviours, accuracy was relatively low (<75%) for all methods, as some specific behaviours were very poorly predicted (e.g. range of F1-scores for vigilance [0-29] % for both sensor types). However, behaviours that were poorly predicted were often confused with other behaviours of the same behavioural class (e.g. vigilance predicted as lying down or standing still, see Tables S4 a.b.). When considering the five behavioural classes, the RF and SVM algorithms performed best, and all of the behavioural classes were predicted with a high level of accuracy for calibration (Table S3), or with a higher level of accuracy than other approaches for validation (Table S8). Indeed, when using the CART and ANN algorithms, some behavioural classes were very poorly predicted (ex: F1 score for grooming =0 for sensor type B, Table S10). Therefore, we selected the RF algorithm for subsequent analyses, as it performed the best, particularly during validation, and needed less computation time than SVM. Indeed, this approach has been widely used for behavioural prediction (e.g. in Eurasian beavers (Graf *et al*. 2015), in puma (Wang *et al*. 2015), in grey seals (Shuert *et al*. 2018)), facilitating comparison among studies. It should be noted that the accelerometer variables used in the RF algorithm did not contribute equally to the discrimination of the 25 behaviours, as their importance varied between 0 and 100% (Appendices S6 & S7).

#### 1) Calibration: Algorithm performance on data from captive animals using Random Forests

RF algorithms were able to predict the 25 behaviours with an accuracy of 56% (type A sensor) and 67% (type B sensor). Classification success differed greatly among behaviours (Fig. 2 and Tables S4 a.b.): while some behaviours were very well predicted (e.g. F1 scores for galloping = 91 % for both sensor types), others were poorly so (e.g. F1 scores for ruminating with head up = 25% and 39% for type A and B, respectively). However, when considering the five behavioural classes (Fig. 2 and Appendix S5), accuracy was 90% and 93% for type A and B sensors, respectively. In particular, foraging head down, walking and unmoving were very well predicted (F1-scores between 84 and 96%, Fig. 2). Running and grooming were slightly less well predicted, especially for type A sensors (F1-scores: 72% and 67% for type A sensor, and 93% and 78% for type B sensor).

**Figure 2:**
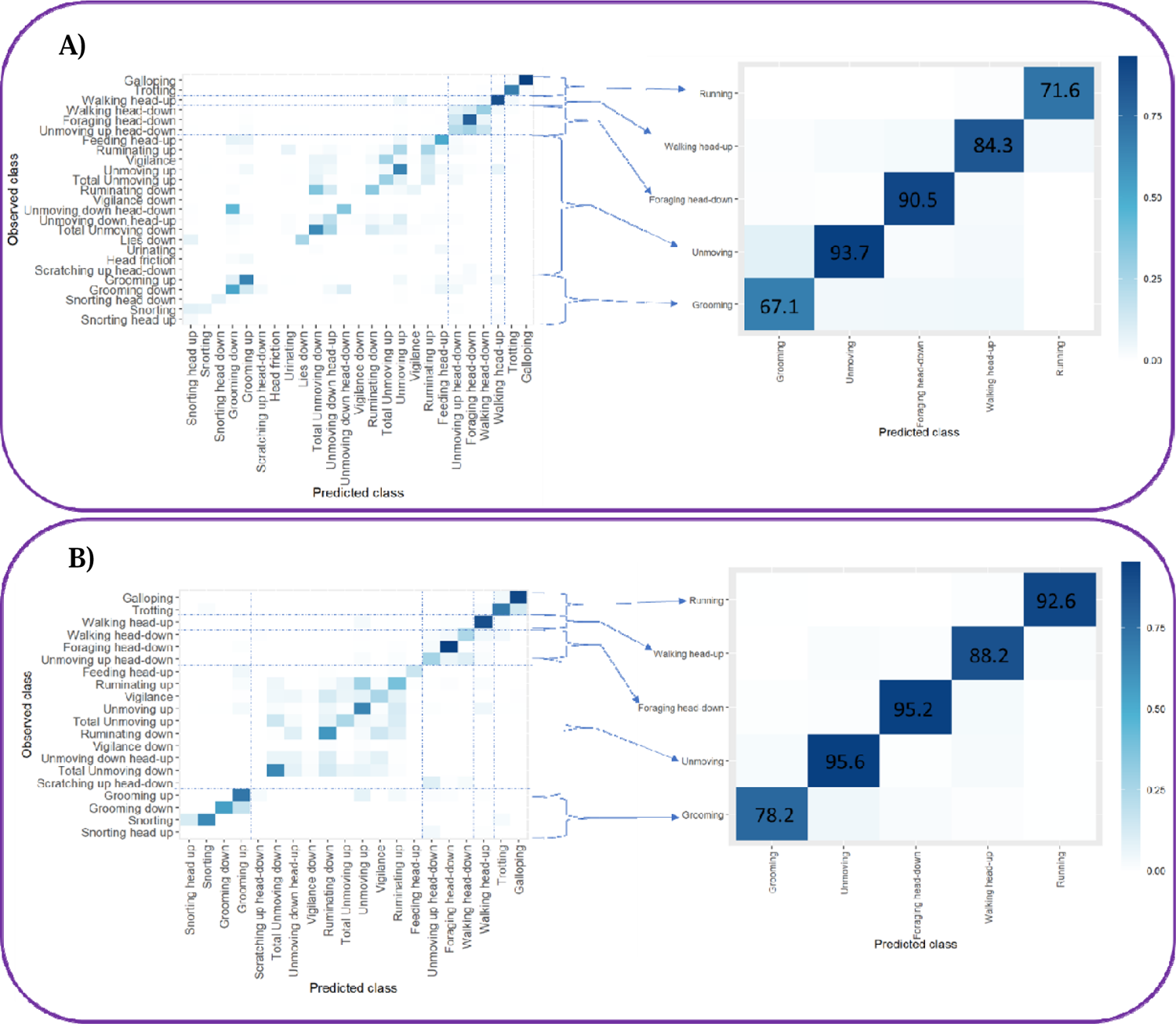
Confusion matrix for the 25 behaviours (left) and the five behavioural classes (right) for (A) the accelerometer type A sensor (N = 3 individuals) and (B) the accelerometer type B sensor (N = 2 individuals). Cross-validation comparing observed vs predicted behaviours was obtained with the Random Forest algorithm applied to the test dataset on captive females. Results are shown as F1 scores for each class (%).

#### 2) Validation: algorithm performance on data from free-ranging animals using Random Forests

Global RF algorithms predicted the five behavioural classes with an accuracy of 90% (range of 86-97% among animals) and 85% (range of 82-91% among animals) for type A and B sensors, respectively (a comparison of predicted and observed behavioural classes for an observed acceleration sequence is provided in Appendix S11).

The five behavioural classes were not predicted with similar accuracy, and the level of accuracy differed slightly from the calibration step (see Fig. 3 and Appendices S5 & S10). The majority of behavioural classes were slightly less well predicted during the validation step (e.g. for unmoving: 94% vs 94 % for sensor type A and 96% vs 92 % for sensor type B), but this difference was more marked for grooming (67% vs 38% for sensor type A and 78% vs 34 % for sensor type B). Thus, global classification algorithms performed very well for identifying foraging head down (F1-scores of 92% and 86% for sensor types A and B, respectively), running (90% and 81% for sensor types A and B, respectively) and unmoving (94% and 92 % for sensor types A and B, respectively) of free-ranging roe deer, but a little less well for walking (83% and 67% for sensor types A and B, respectively), and rather badly for grooming (38% and 34% for sensor types A and B, respectively). The F1-score values were close to those for both sensitivity and precision. This means that no behaviour was more overpredicted (lower values of precision) than underpredicted (lower values of precision), or vis a versa. Overall, for both sensor types, most of the misclassified values for grooming were confused with foraging head-down or unmoving, whereas for sensor type B only, misclassified values for walking were mostly confused with unmoving.

**Figure 3:**
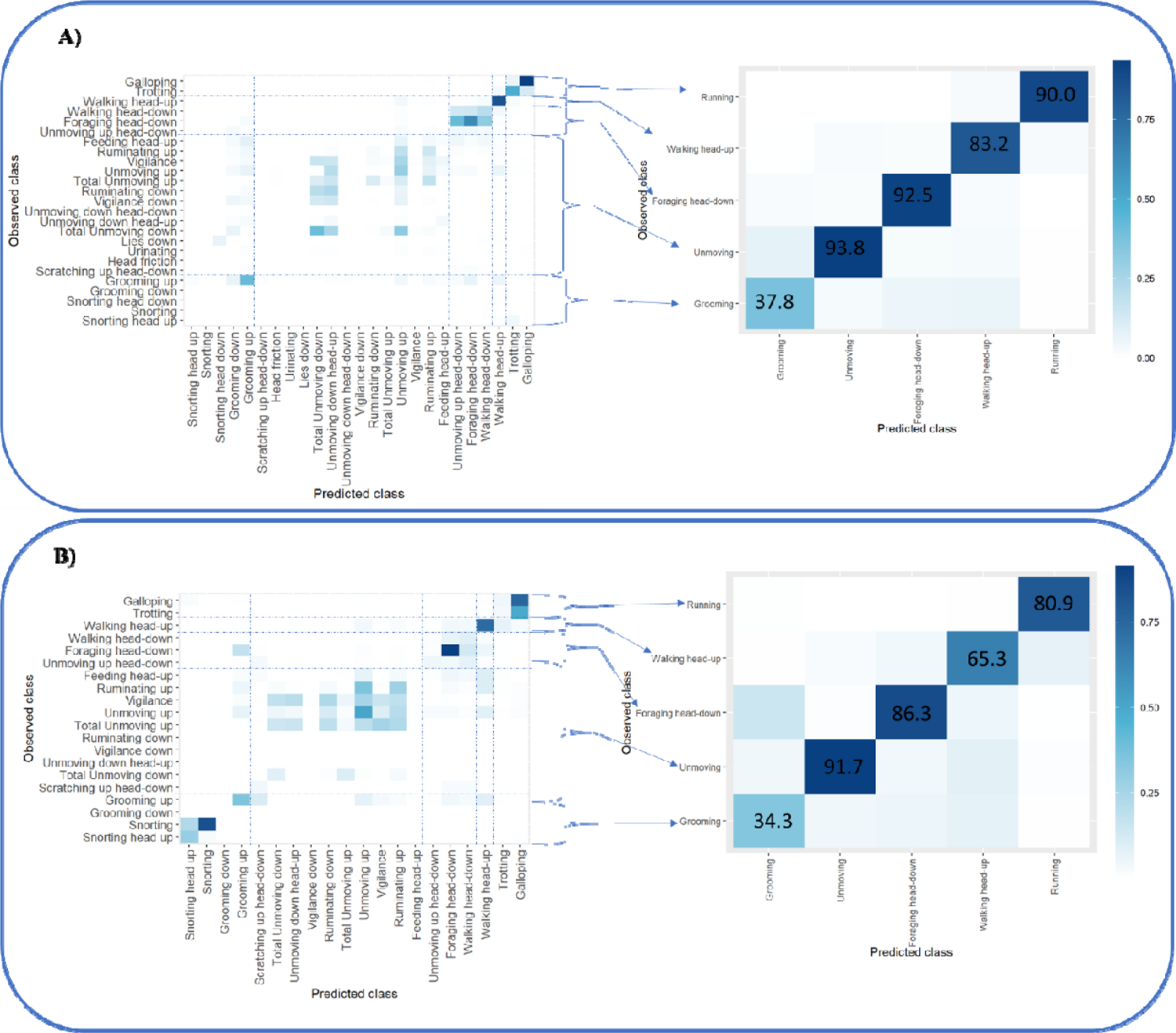
Confusion matrix for the 25 behaviours and the five behavioural classes for (A) the sensor type A (N = 6 individuals) and (B) the sensor type B (N = 8 individuals). Cross-validation of observed versus predicted classes was obtained with the Random Forest algorithm applied to the test dataset of the free-ranging roe deer. Results are shown as F1 scores for each class (%).

#### 3) Application: Algorithm predictions for the circadian behavioural rhythm of free-ranging roe deer

We estimated the circadian time-budget of free-ranging roe deer during March with respect to the five behavioural classes using the predictions of the global classification algorithms. First, according to the Bayesian multilevel mixed model describing variation in the proportion of time allocated to each behavioural class per hour, we found that free-ranging roe deer spent most of their active time foraging head down, i.e. on average, 4.4±0.6 h per day, followed by 2.0±0.5 h walking, 1.8±0.4 h grooming, and 0.2±0.1 h running. The remaining time was allocated to unmoving behaviours other than grooming, including standing, lying down, vigilance or feeding with head-up, representing around 65% of the total time (15.7±0.8 h) (Fig. 4). Hence, roe deer were actually engaged in foraging activities for two-thirds (on average 4.4 hours) of their habitual time active (total of 6.6 hours per day, either foraging, walking or running). This time-budget, however, varied greatly over the day. Indeed, the selected model that best described variation in the proportion of time allocated to each behavioural class per hour included a smoothed effect of the time of day (Delta LOOIC with the basic model: 12790.1), such that deer were more active during twilight hours (defined here as two hours around either sunrise or sunset). During March, they spent around twice as much time foraging head-down, walking or running during dawn and dusk than during the day or night (Fig. 5.a,b,c) (Fig. 5.d). More precisely, we observed that foraging occurred around half as frequently during the day than during other periods (mean percentage of time foraging head down during the day: 7.9±1.7 % vs 16.2±3.1 % during other periods), whereas moving behaviours were clearly concentrated during twilight (mean percentage of time walking or running during twilight: 13.6±1.5 % and 3.1±0.6 %, vs. during day and night: 7.5±1.0 % and 1.9±0.4 %). Only the time allocated to grooming behaviour did not vary over the course of the day (Fig. 5.e).

**Figure 4:**
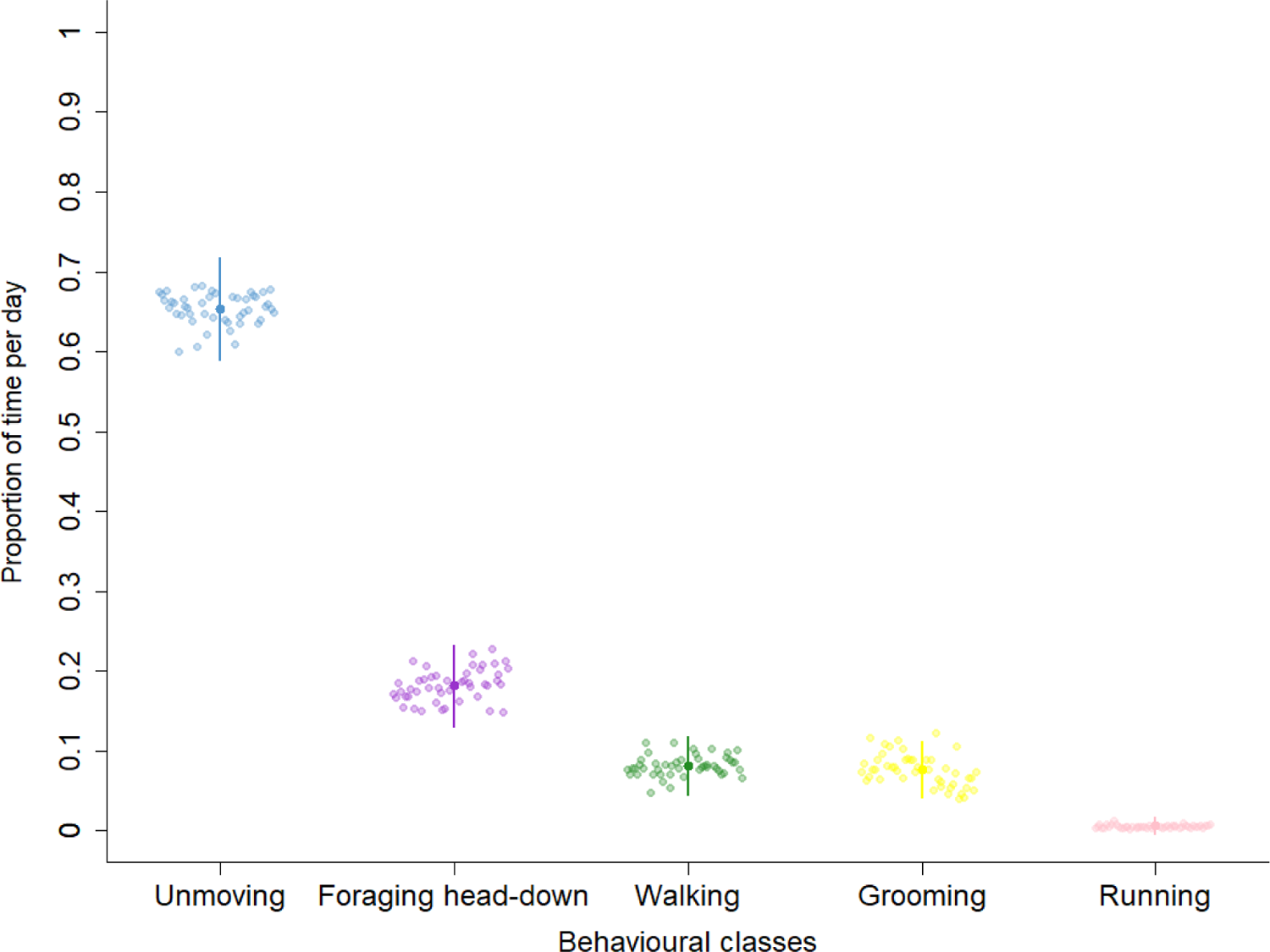
Daily time budget of 47 free-ranging roe deer during March based on the five behavioural classes inferred with the global classification algorithm. Solid points (and bars) represent predictions (and their associated 95% credible intervals). Hollow points represent observed values averaged for one individual.

**Figure 5:**
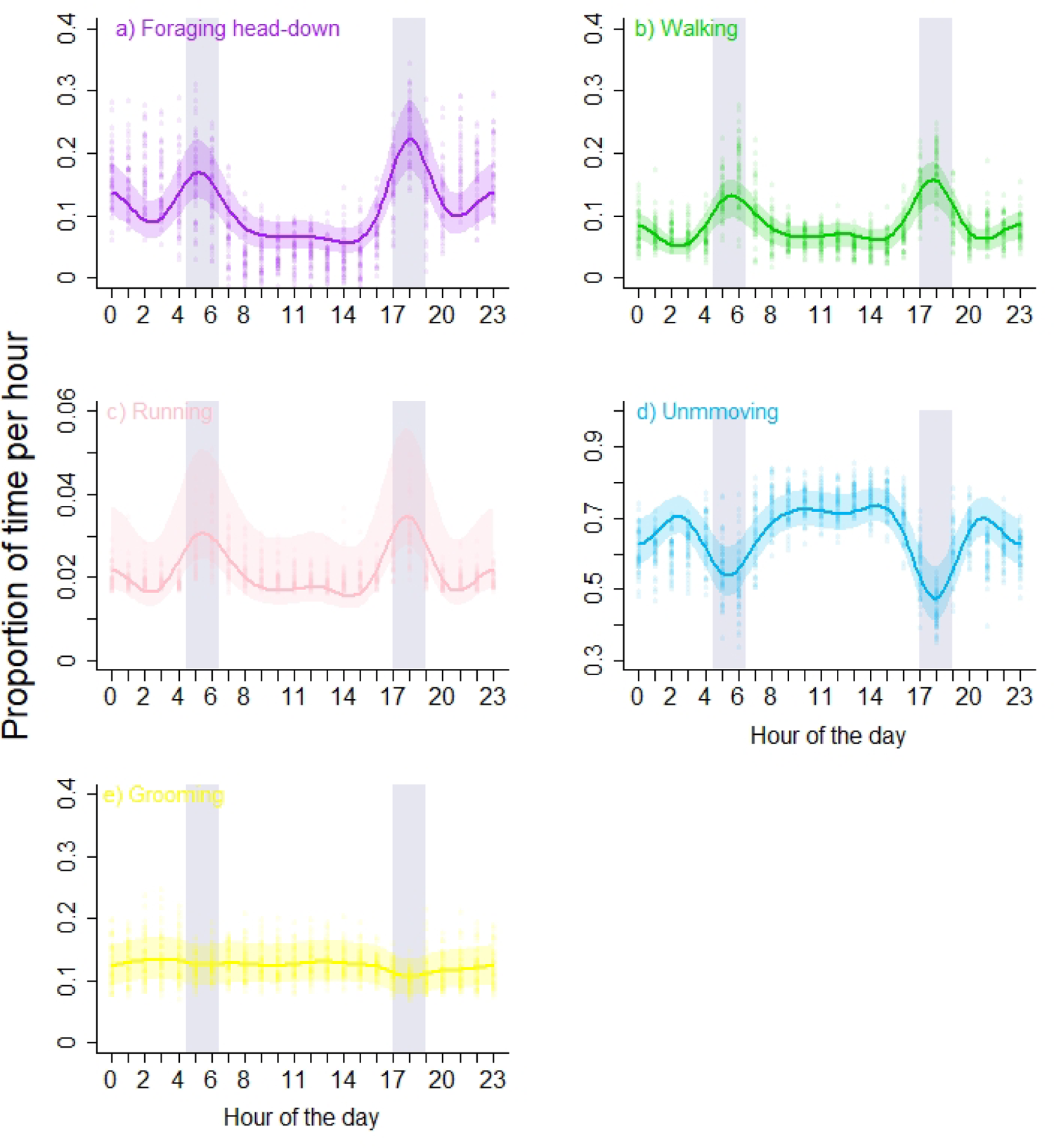
Hourly time budget for 47 free-ranging roe deer based on the five behavioural classes (a-e) inferred with the global classification algorithm. Solid lines (and the coloured area) represent predictions (and their associated 95% credible intervals). Points represent observed values averaged for one individual. Grey bars indicate two hours around mean twilight during March. Hours of the day are in GMT.

## Discussion

The golden age of biologging (Wilmers *et al*. 2015) is revolutionising our capacity to understand the behavioural ecology of wild animals. Using high frequency accelerometer monitoring, we successfully identified the most important behavioural phases of wild free-ranging cryptic roe deer with a high degree of accuracy (> 85%). We obtained daily and hourly time budgets of 47 free-ranging individuals, providing evidence that such detailed information may help to better understand the behaviour of this species in relation to environmental variability. The calibration and validation procedures that we developed are equally applicable to other species, as long as high quality accelerometric and behavioural data are available to build a sufficiently powerful classification algorithm. Below, we discuss the strengths and limitations of our method, and provide guidelines (also summarized in Box 1) and potential future directions for elucidating the behaviour of wild animals with this approach.

### Calibration of a classification algorithm with captive animals

The calibration procedure using captive roe deer enabled us to predict discrete behavioural categories, specifically, walking, running, foraging head-down, grooming and unmoving, with a high overall level of accuracy. Furthermore, using a classification algorithm based on a detailed ethogram of 25 specific behaviours, we were able to reliably identify certain specific behaviours (e.g. F1-scores > 90% for galloping), which could be the target of specific research questions. For example, accurate predictions for the occurrence of galloping may provide insight on flight behaviour in response to human disturbance (Stankowich 2008b), thus providing a better description of the landscape of fear (Laundre *et al*. 2010) for wild, cryptic species.

We used captive animals for the calibration step for their ease of observation (6 to 10 hours of film per sensor type), generating a large dataset of acceleration signals and associated behaviours. However, certain behaviours did not figure in our classification algorithm because they were rarely or never observed in captivity (e.g. fighting, mating and nursing), although accelerometers are powerful tools to detect these rare events (e.g. mating behaviour in sharks (*Ginglymostoma cirratum*) (Whitney *et al*. 2010); display behaviours in little bustards (*Tetrax tetrax*) (Gudka *et al*. 2019)). For behaviours such as parturition (Fogarty *et al*. 2020; Gurule *et al*. 2021) or maternal care (ex: in grey seal (*Halichoerus grypus*) Shuert *et al*. 2020) that are difficult to observe in the wild, it is, therefore, key to ensure that captive animals are able to express them. The advantages and limitations of relying on captive animals to calibrate behaviour classification algorithms needs to be carefully considered on a case by case basis (***Guideline 1***).

A crucial parameter for algorithm performance is the position of the accelerometer on the animal’s body (Barwick *et al*. 2018, ***Guideline 2***). For many species, accelerometers are fixed on a collar which is positioned on the animal’s neck, which may impair detection of important behaviours that do not involve marked movements of the head (Tables S4). For example, in our analysis, foraging with head up was poorly discriminated (F1-scores < 50%), probably because the associated movements were not distinguishable from unmoving behaviours such as observation or vigilance. Studies on livestock have used accelerometers located near the jaw to detect chewing (Alvarenga *et al*. 2016), but this is currently difficult to use on wild animals, unless a suitable system is developed that does not significantly impact animal welfare. Alternatively, foraging and rumination may be more efficiently detected by combining accelerometery with an acoustic sensor (Studd *et al*. 2019a)

Because biologging is a young and rapidly developing field, manufacturers are constantly upgrading their equipment (Boyd *et al*. 2004; Evans *et al*. 2013). As was the case in this study, ecological studies often span years, or even decades (Lindenmayer *et al*. 2012), so that calibration must account for technological developments (***Guideline 3***). Indeed, despite using a single manufacturer, we found that the acceleration signal varied according to the orientation of the axes of a particular sensor, sampling frequency and tag position on the animals’ neck due to weight differences between versions of the sensor. As a result, key parameters of the procedure (see Fig. 1) must be established independently for each type of equipment, as we did for the two sensor types (e.g. number of feature variables, time window duration to calculate variables derived from raw acceleration data, choice of machine learning approach).

Finally, accelerometery generates huge data sets, and there is a trade-off between the precision of behavioural predictions and the time and space needed for storage and to process the data. In this study, we found little difference in classification performance in relation to sampling frequency (8 or 32 Hz, see example in Table S3), suggesting that lower rates of data acquisition sufficed for our purposes (see Hounslow *et al*. 2019). As in several studies using accelerometer data (Fehlmann *et al*. 2017; Tatler *et al*. 2018), we included a wide diversity of accelerometer-related variables in the algorithms (see Table 2), which improved their predictive power, but also significantly increased computation time (Fig. 1 - Part 2). In our case, RF and SVM provided the most powerful algorithms, but were also the longest to implement in terms of computing time, algorithm construction (RF) and prediction (SVM) (Fig. 1 - Part 3&4). The choice of method will, thus, depend on the target behaviour studied and, more generally, the trade-off between precision and effort to process the accelerometer big data (***Guideline 4***).

### Validating classification algorithms for estimating behaviour in the wild

The use of free-ranging roe deer to validate the RF algorithms that we built based on captive roe deer confirmed the efficiency of the algorithms for predicting the main behavioural classes that constitute the activity of this species (accuracy> 85%). Indeed, except for grooming behaviours (F1-scores < 40%), the RF algorithms were able to predict foraging head down, running, walking and unmoving with high performance in free-ranging roe deer (range of F1-scores: [83-94%] for sensor type A and [68-92%] for sensor type B).

This validation step using several roe deer allowed us to identify that acceleration signals vary among individuals in relation to collar tightness and sensitivity (see section on accelerometer characteristics) which can influence the performance of the algorithms. It is, hence, important to take these variations into account for behavioural prediction across a number of individuals (***Guideline 5***). Problems of sensor variability could be resolved by applying data correction, if manufacturers provide information to control this source of variability (Kröschel *et al*. 2017), or by using acceleration variables which summarize acceleration over the three axes, such as VeDBA and OBDA (Studd *et al*. 2019b). Another solution, that we used in this study, is to scale the data per individual, which provided better behavioural prediction for wild roe deer. Despite these procedures, differences in performance between supervised machine learning methods were more marked for the validation step (difference in accuracy between the best and worst methods: 11 % for sensor type A and 28 % for sensor type B) than for the calibration step (difference in accuracy between the best and worst methods: 6 % for sensor type A and 8 % for sensor type B), and, globally, algorithm performance tended to decrease between steps. We think that this could be due to residual variation in the data across individuals and sensors that we could not control for. However, it could also be because we were unable to record the full range of behaviours that the classification algorithm can predict with the quite small number of videos that we obtained per free-ranging individual (see Appendix S2),. Thus, we consider that it is important to validate the classification algorithms on independent individuals, which can alter the performance of the model predictions (Ferdinandy *et al*. 2020), and particularly on free-ranging individuals in order to estimate the performance of the algorithms on data similar to those used in the application step (***Guideline 6***).

### Inferring the circadian time budget of free-ranging animals

The use of accelerometers to infer the time-budget of free-ranging roe deer is a promising approach, as has been demonstrated in other species such as red foxes (*Vulpes vulpes*) (Rast et al. 2020), snowshoe hares (*Lepus americanus*) (Studd et al. 2019a), fur seals (*Arctocephalus pusillus*) (Ladds et al. 2018), chipmunks (*Tamias alpinus & T. speciosus*) (Hammond et al. 2016) and crab plovers (*Dromas ardeola*) (Bom et al. 2014). For roe deer, previous studies have often been limited to information on space use (Cagnacci *et al*. 2010; Malagnino *et al*. 2021) and activity (Kozakai *et al*. 2013; Pagon *et al*. 2013), or to occasional direct observations of specific behaviours (e.g. vigilance (Benhaiem *et al*. 2008; Favreau *et al*. 2014)), usually during the day and in habitats that provide high observability. Using an approach based on accelerometer sensors, ecologists will be increasingly able to remotely and continuously estimate specific behaviours of wild animals and how they may vary among individuals (Bidder *et al*. 2020; Bennison *et al*. 2022) and over time (e.g. according time of day and season (Hammond *et al*. 2016; Rast *et al*. 2020), according to environmental conditions).

Here, using continuous accelerometer data obtained on 47 free-ranging roe deer, we were able to infer the behavioural time budget across the 24-hour diel cycle, irrespective of their habitat or location. For example, we were able to identify foraging head-down behaviour with a high level of accuracy, and showed that free-ranging roe deer spent, on average, more than four hours per day engaged in this activity. However, this was particularly concentrated during twilight (Fig. 4), but much less prevalent during daylight, presumably due to frequent disturbance in this human-dominated area (Bonnot *et al*. 2013). Interestingly, we found that roe deer also spent much more time moving during twilight, which may reflect the constraints of travelling from refuge areas frequented during daylight to richer feeding grounds in highly fragmented landscapes (Abbas *et al*. 2011). However, it should be noted that the classification algorithms struggled to accurately identify foraging with head up, a common behaviour in browsers such as roe deer, at least during certain seasons (Bodmer 1990). Indeed, during spring, roe deer are known to feed on buds and young leaves of trees and shrubs, when available (Tixier & Duncan 1996), so that we almost certainly underestimated total foraging time. It is hence important to bear in mind the limitations of a given classification algorithm when interpreting the results (***Guideline 7***).

## Conclusion and perspectives

The approach that we present in this paper, combining calibration and validation of the accelerometer prior to application in a free-ranging animal, is transferable to other species and will enable accurate prediction of important behaviours in cryptic wild animals. Our practical guidelines for improving behavioural prediction highlight the importance of the validation step which is frequently omitted (Wang *et al*. 2015; Ladds *et al*. 2018; Rast *et al*. 2020). Indeed, validation through field verification on free-ranging individuals is crucial to ensure robust behavioural inference. The combination of accelerometers with GPS monitoring opens the door to spatially explicit modelling of behaviour (Wang *et al*. 2015; Rast *et al*. 2020; Tatler *et al*. 2021), for example, to better understand the impact of landscape modifications on wild populations. Similarly, combining accelerometers with acoustic sensors could aid identification of discrete behaviours such as chewing, vocalization or grooming (Wijers *et al*. 2018, 2021; Studd *et al*. 2019a), for example, to better document the responses of wildlife to human disturbance or predation risk (Lynch *et al*. 2015). In the context of the ever-increasing anthropogenic pressure, biologging promises to help us better understand how wildlife copes with human-induced rapid environmental change.

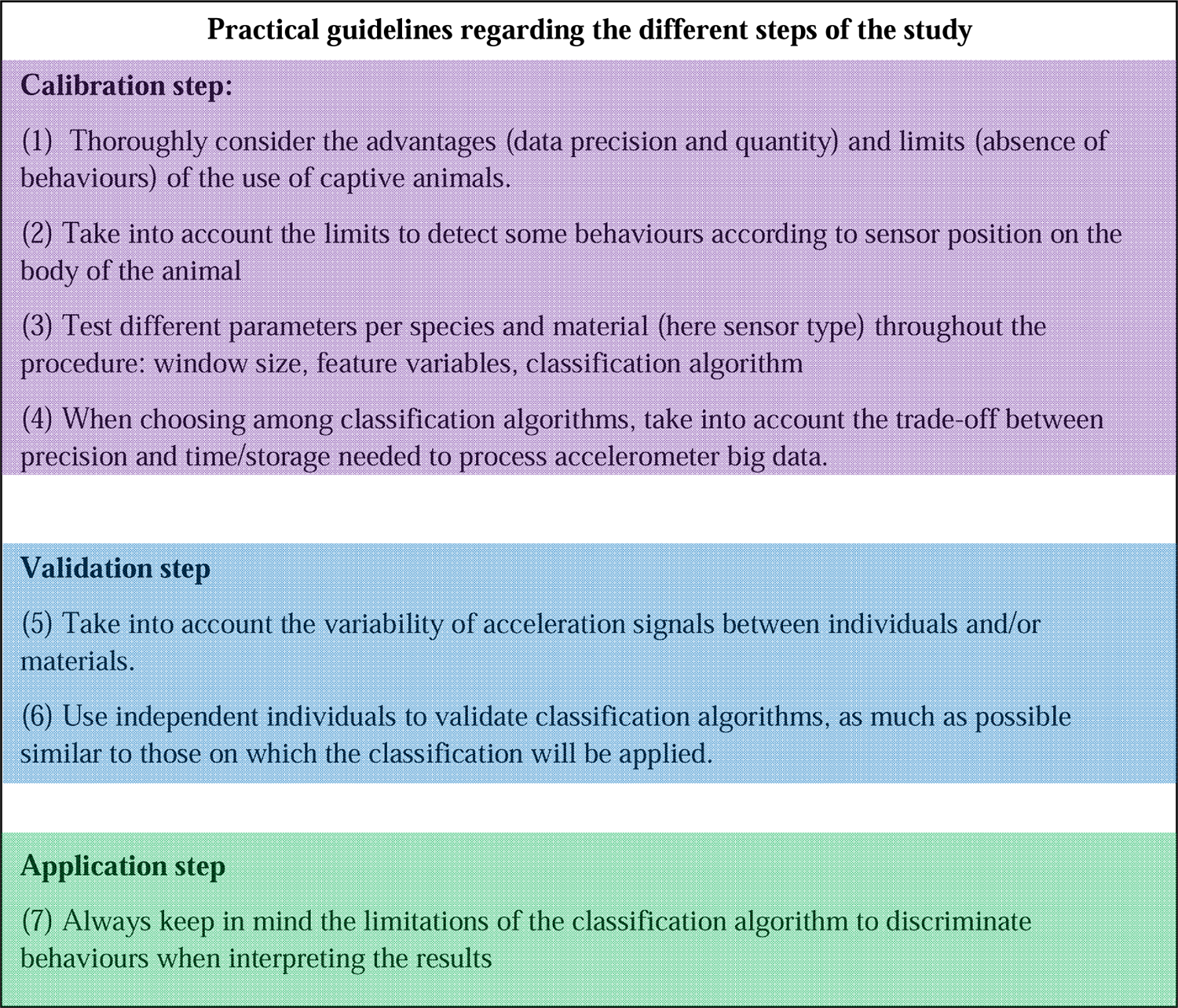

## References

Abbas, F., Morellet, N., Hewison, A.J.M., Merlet, J., Cargnelutti, B., Lourtet, B., et al. (2011). Landscape fragmentation generates spatial variation of diet composition and quality in a generalist herbivore. Oecologia, 167, 401–411.

Alvarenga, F.A.P., Borges, I., Palkovič, L., Rodina, J., Oddy, V.H. & Dobos, R.C. (2016). Using a three-axis accelerometer to identify and classify sheep behaviour at pasture. Applied Animal Behaviour Science, 181, 91–99.

Barwick, J., Lamb, D.W., Dobos, R., Welch, M. & Trotter, M. (2018). Categorising sheep activity using a tri-axial accelerometer. Computers and Electronics in Agriculture, 145, 289– 297.

Benhaiem, S., Delon, M., Lourtet, B., Cargnelutti, B., Aulagnier, S., Hewison, A.J.M., et al. (2008). Hunting increases vigilance levels in roe deer and modifies feeding site selection. Animal Behaviour, 76, 611–618.

Bennison, A., Giménez, J., Quinn, J., Green, J. & Jessopp, M. (2022). A bioenergetics approach to understanding sex differences in the foraging behaviour of a sexually monomorphic species. Royal Society Open Science, 9.

Bidder, O., di Virgilio, A., Hunter, J., Mcinturff, A., Gaynor, K., Smith, A., et al. (2020). Monitoring canid scent marking in space and time using a biologging and machine learning approach. Scientific Reports, 10, 588.

Bidder, O.R., Walker, J.S., Jones, M.W., Holton, M.D., Urge, P., Scantlebury, D.M., et al. (2015). Step by step: reconstruction of terrestrial animal movement paths by dead-reckoning. Movement Ecology, 3.

Bodmer, R.E. (1990). Ungulate Frugivores and the Browser-Grazer Continuum. Oikos, 57, 319.

Bom, R.A., Bouten, W., Piersma, T., Oosterbeek, K. & van Gils, J.A. (2014). Optimizing acceleration-based ethograms: the use of variable-time versus fixed-time segmentation. Movement Ecology, 2.

Bonnot, N., Morellet, N., Verheyden, H., Cargnelutti, B., Lourtet, B., Klein, F., et al. (2013). Habitat use under predation risk: hunting, roads and human dwellings influence the spatial behaviour of roe deer. European Journal of Wildlife Research, 59, 185–193.

Bonnot, N.C., Couriot, O., Berger, A., Cagnacci, F., Ciuti, S., Groeve, J.E.D., et al. (2020). Fear of the dark? Contrasting impacts of humans versus lynx on diel activity of roe deer across Europe. Journal of Animal Ecology, 89, 132–145.

Bonnot, N.C., Hewison, A.J.M., Morellet, N., Gaillard, J.-M., Debeffe, L., Couriot, O., et al. (2017). Stick or twist: roe deer adjust their flight behaviour to the perceived trade-off between risk and reward. Animal Behaviour, 124, 35–46.

Boyd, I.L., Kato, A. & Ropert-Coudert, Y. (2004). Bio-logging science: sensing beyond the boundaries, 15.

Breiman, L. (2001). Random Forests. Machine Learning, 45, 5–32.

Brown, D.D., Kays, R., Wikelski, M., Wilson, R. & Klimley, A. (2013). Observing the unwatchable through acceleration logging of animal behavior. Animal Biotelemetry, 1, 20.

Bürkner, P.-C. (2017). brms: An R package for Bayesian multilevel models using Stan. Journal of statistical software, 80, 1–28.

Cagnacci, F., Boitani, L., Powell, R.A. & Boyce, M.S. (2010). Animal ecology meets GPS-based radiotelemetry: a perfect storm of opportunities and challenges. Philosophical Transactions of the Royal Society B: Biological Sciences, 365, 2157–2162.

Caro, T. (1999). The behaviour–conservation interface. Trends in ecology & evolution, 14, 366–369.

Chicco, D. & Jurman, G. (2020). The advantages of the Matthews correlation coefficient (MCC) over F1 score and accuracy in binary classification evaluation. BMC Genomics, 21, 6.

Chicco, D., Starovoitov, V. & Jurman, G. (2021). The Benefits of the Matthews Correlation Coefficient (MCC) Over the Diagnostic Odds Ratio (DOR) in Binary Classification Assessment. *IEEE Acces*s, PP, 1–1.

Collins, P.M., Green, J.A., Warwick-Evans, V., Dodd, S., Shaw, P.J.A., Arnould, J.P.Y., et al. (2015). Interpreting behaviors from accelerometry: a method combining simplicity and objectivity. Ecology and Evolution, 5, 4642–4654.

Cooke, S.J., Hinch, S.G., Wikelski, M., Andrews, R.D., Kuchel, L.J., Wolcott, T.G., et al. (2004). Biotelemetry: a mechanistic approach to ecology. Trends in Ecology & Evolution, 19, 334–343.

Cozzi, G., Behr, D.M., Webster, H.S., Claase, M., Bryce, C.M., Modise, B., et al. (2020). African Wild Dog Dispersal and Implications for Management. The Journal of Wildlife Management.

Dickinson, E.R., Stephens, P.A., Marks, N.J., Wilson, R.P. & Scantlebury, D.M. (2020). Best practice for collar deployment of tri-axial accelerometers on a terrestrial quadruped to provide accurate measurement of body acceleration. Animal Biotelemetry, 8.

Ditmer, M.A., Rettler, S.J., Fieberg, J.R., Iaizzo, P.A., Laske, T.G., Noyce, K.V., et al. (2018). American black bears perceive the risks of crossing roads. Behavioral Ecology, 29, 667–675.

Dujon, A.M., Schofield, G., Lester, R.E., Esteban, N. & Hays, G.C. (2017). Fastloc-GPS reveals daytime departure and arrival during long-distance migration and the use of different resting strategies in sea turtles. Mar Biol, 164, 187.

Evans, K., Lea, M.-A. & Patterson, T.A. (2013). Recent advances in bio-logging science: Technologies and methods for understanding animal behaviour and physiology and their environments. Deep Sea Research Part II: Topical Studies in Oceanography, 88–89, 1–6.

Favreau, F.-R., Goldizen, A.W., Fritz, H., Blomberg, S.P., Best, E.C. & Pays, O. (2014). Within-population differences in personality and plasticity in the trade-off between vigilance and foraging in kangaroos. Animal Behaviour, 92, 175–184.

Fehlmann, G., O’Riain, M.J., Hopkins, P.W., O’Sullivan, J., Holton, M.D., Shepard, E.L.C., et al. (2017). Identification of behaviours from accelerometer data in a wild social primate. Animal Biotelemetry, 5.

Ferdinandy, B., Gerencsér, L., Corrieri, L., Perez, P., Újváry, D., Csizmadia, G., et al. (2020). Challenges of machine learning model validation using correlated behaviour data: Evaluation of cross-validation strategies and accuracy measures. PLOS ONE, 15, e0236092.

Fogarty, E.S., Swain, D.L., Cronin, G.M., Moraes, L.E. & Trotter, M. (2020). Can accelerometer ear tags identify behavioural changes in sheep associated with parturition? Animal Reproduction Science, 216, 106345.

Friard, O. & Gamba, M. (2016). BORIS: a free, versatile open-source event-logging software for video/audio coding and live observations. Methods in Ecology and Evolution, 7, 1325– 1330.

Gleiss, A.C., Wilson, R.P. & Shepard, E.L.C. (2011). Making overall dynamic body acceleration work: on the theory of acceleration as a proxy for energy expenditure: Acceleration as a proxy for energy expenditure. Methods in Ecology and Evolution, 2, 23–33.

Graf, P.M., Wilson, R.P., Qasem, L., Hackländer, K. & Rosell, F. (2015). The Use of Acceleration to Code for Animal Behaviours; A Case Study in Free-Ranging Eurasian Beavers Castor fiber. PLOS ONE, 10, e0136751.

Grémillet, D., Kuntz, G., Woakes, A., Gilbert, C., Robin, J., Le Maho, Y., et al. (2005). Year-round recording of behavioural and physiological parameters reveal the survival strategy of a poorly insulated diving endotherm during the Arctic winter. The Journal of experimental biology, 208, 4231–41.

Grémillet, D., Lescroël, A., Ballard, G., Dugger, K.M., Massaro, M., Porzig, E.L., et al. (2018). Energetic fitness: Field metabolic rates assessed via 3D accelerometry complement conventional fitness metrics. Functional Ecology, 32, 1203–1213.

Gudka, M., Santos, C.D., Dolman, P.M., Abad-Gómez, J.M. & Silva, J.P. (2019). Feeling the heat: Elevated temperature affects male display activity of a lekking grassland bird. PLOS ONE, 14, e0221999.

Gurarie, E., Bracis, C., Delgado, M., Meckley, T.D., Kojola, I. & Wagner, C.M. (2016). What is the animal doing? Tools for exploring behavioural structure in animal movements. Journal of Animal Ecology, 85, 69–84.

Gurule, S.C., Tobin, C.T., Bailey, D.W. & Hernandez Gifford, J.A. (2021). Evaluation of the tri-axial accelerometer to identify and predict parturition-related activities of Debouillet ewes in an intensive setting. Applied Animal Behaviour Science, 237, 105296.

Hammond, T.T., Springthorpe, D., Walsh, R.E. & Berg-Kirkpatrick, T. (2016). Using accelerometers to remotely and automatically characterize behavior in small animals. The Journal of Experimental Biology, 219, 1618–1624.

Hewison, A.J.M, Vincent, J., Joachim, J., Angibault, J., Cargnelutti, B. & Cibien, C. (2001). The effects of woodland fragmentation and human activity on roe deer distribution in agricultural landscapes. Canadian Journal of Zoology, 79, 679–689.

Hewison, A.J.M., Morellet, N., Verheyden, H., Daufresne, T., Angibault, J.-M., Cargnelutti, B., et al. (2009). Landscape fragmentation influences winter body mass of roe deer. Ecography, 32, 1062–1070.

Hounslow, J.L., Brewster, L.R., Lear, K.O., Guttridge, T.L., Daly, R., Whitney, N.M., et al. (2019). Assessing the effects of sampling frequency on behavioural classification of accelerometer data. Journal of Experimental Marine Biology and Ecology, 512, 22–30.

Kooyman, G.L. (2004). Genesis and evolution of bio-logging devices: 1963–2002, 8.

Kozakai, C., Yamazaki, K., Nemoto, Y., Nakajima, A., Umemura, Y., Koike, S., et al. (2013). Fluctuation of daily activity time budgets of Japanese black bears: relationship to sex, reproductive status, and hard-mast availability. Journal of Mammalogy, 94, 351–360.

Krop-Benesch, A., Berger, A., Hofer, H. & Heurich, M. (2013). Long-term measurement of roe deer (Capreolus capreolus) (Mammalia: Cervidae) activity using two-axis accelerometers in GPS-collars. Italian Journal of Zoology, 80, 69–81.

Kröschel, M., Reineking, B., Werwie, F., Wildi, F. & Storch, I. (2017). Remote monitoring of vigilance behavior in large herbivores using acceleration data. Animal Biotelemetry, 5.

Ladds, M.A., Salton, M., Hocking, D.P., McIntosh, R.R., Thompson, A.P., Slip, D.J., et al. (2018). Using accelerometers to develop time-energy budgets of wild fur seals from captive surrogates. PeerJ, 6, e5814.

Laundre, J.W., Hernandez, L. & Ripple, W.J. (2010). The Landscape of Fear: Ecological Implications of Being Afraid∼!2009-09-09∼!2009-11-16∼!2010-02-02∼! The Open Ecology Journal, 3, 1–7.

Lindenmayer, D.B., Likens, G.E., Andersen, A., Bowman, D., Bull, C.M., Burns, E., et al. (2012). Value of long-term ecological studies. Austral Ecology, 37, 745–757.

Löttker, P., Rummel, A., Traube, M., Stache, A., Šustr, P., Müller, J., et al. (2009). New Possibilities of Observing Animal Behaviour from a Distance Using Activity Sensors in Gps-Collars: An Attempt to Calibrate Remotely Collected Activity Data with Direct Behavioural Observations in Red Deer *Cervus elaphus*. Wildlife Biology, 15, 425–434.

Lush, L., Ellwood, S., Markham, A., Ward, A.I. & Wheeler, P. (2016). Use of tri-axial accelerometers to assess terrestrial mammal behaviour in the wild. Journal of Zoology, 298, 257–265.

Lynch, E., Northrup, J.M., McKenna, M.F., Anderson, C.R., Angeloni, L. & Wittemyer, G. (2015). Landscape and anthropogenic features influence the use of auditory vigilance by mule deer. Behavioral Ecology, 26, 75–82.

Malagnino, A., Marchand, P., Garel, M., Cargnelutti, B., Itty, C., Chaval, Y., et al. (2021). Do reproductive constraints or experience drive age-dependent space use in two large herbivores? Animal Behaviour, 172, 121–133.

Marchand, P., Garel, M., Morellet, N., Benoit, L., Chaval, Y., Itty, C., et al. (2021a). A standardised biologging approach to infer parturition: An application in large herbivores across the hider-follower continuum. Methods in Ecology and Evolution, 12, 1017–1030.

Marchand, T., Gal, A.-S.L. & Georges, J.-Y. (2021b). Fine scale behaviour and time-budget in the cryptic ectotherm European pond turtle *Emys orbicularis*. PLOS ONE, 16, e0256549.

Montané, J., Marco, I., López-Olvera, J., Perpiñán, D., Manteca, X. & Lavín, S. (2003). Effects of acepromazine on capture stress in roe deer (*Capreolus capreolus*). Journal of Wildlife Diseases, 39, 375–386.

Morellet, N., Van Moorter, B., Cargnelutti, B., Angibault, J.-M., Lourtet, B., Merlet, J., et al. (2011). Landscape composition influences roe deer habitat selection at both home range and landscape scales. Landscape Ecology, 26, 999–1010.

Mosser, A.A., Avgar, T., Brown, G.S., Walker, C.S. & Fryxell, J.M. (2014). Towards an energetic landscape: broad-scale accelerometry in woodland caribou. Journal of Animal Ecology, 83, 916–922.

Nathan, R., Spiegel, O., Fortmann-Roe, S., Harel, R., Wikelski, M. & Getz, W.M. (2012). Using tri-axial acceleration data to identify behavioral modes of free-ranging animals: general concepts and tools illustrated for griffon vultures. Journal of Experimental Biology, 215, 986– 996.

Padié, S., Morellet, N., Hewison, A.J.M., Martin, J.-L., Bonnot, N., Cargnelutti, B., et al. (2015). Roe deer at risk: teasing apart habitat selection and landscape constraints in risk exposure at multiple scales. Oikos, 124, 1536–1546.

Pagano, A., Rode, K., Cutting, A., Owen, M., Jensen, S., Ware, J., et al. (2017). Using tri-axial accelerometers to identify wild polar bear behaviors. Endangered Species Research, 32, 19–33.

Pagon, N., Grignolio, S., Pipia, A., Bongi, P., Bertolucci, C. & Apollonio, M. (2013). Seasonal variation of activity patterns in roe deer in a temperate forested area. Chronobiology International, 30, 772–785.

Ponganis, P.J., Kooyman, G.L., Starke, L.N., Kooyman, C.A. & Kooyman, T.G. (1997). Post-dive blood lactate concentrations in emperor penguins, Aptenodytes forsteri. J Exp Biol, 200, 1623–1626.

Ponganis, P.J., Ponganis, E.P., Ponganis, K.V., Kooyman, G.L., Gentry, R.L. & Trillmich, F. (1990). Swimming velocities in otariids. Can. J. Zool., 68, 2105–2112.

Qasem, L., Cardew, A., Wilson, A., Griffiths, I., Halsey, L.G., Shepard, E.L.C., et al. (2012). Tri-Axial Dynamic Acceleration as a Proxy for Animal Energy Expenditure; Should We Be Summing Values or Calculating the Vector? PLoS ONE, 7, e31187.

Rast, W., Kimmig, S.E., Giese, L. & Berger, A. (2020). Machine learning goes wild: Using data from captive individuals to infer wildlife behaviours. PLOS ONE, 15, e0227317.

R Core Team (2020). R: A language and environment for statistical computing. R Foundation for Statistical Computing, Vienna, Austria. https://www.R-project.org/

Ropert-Coudert, Y., Kato, A., Wilson, R. & Cannell, B. (2006). Foraging strategies and prey encounter rate of free-ranging Little Penguins. Mar. Biol, 149, 139–148.

Sakamoto, K.Q., Sato, K., Ishizuka, M., Watanuki, Y., Takahashi, A., Daunt, F., et al. (2009). Can Ethograms Be Automatically Generated Using Body Acceleration Data from Free-Ranging Birds? PLOS ONE, 4, e5379.

Sato, K. (2003). Factors affecting stroking patterns and body angle in diving Weddell seals under natural conditions. Journal of Experimental Biology, 206, 1461–1470.

Schneirla, T.C. (1950). The relationship between observation and experimentation in the field study of behavior. Annals of the New York Academy of Sciences, 51, 1022–1044.

Seigle-Ferrand, J., Atmeh, K., Gaillard, J.-M., Ronget, V., Morellet, N., Garel, M., et al. (2021). Home range size variation within and across large herbivore populations: what do we know after 50 years of telemetry studies?, Frontiers in Ecology and Evolution, 515.

Shamoun-Baranes, J., Bom, R., van Loon, E.E., Ens, B.J., Oosterbeek, K. & Bouten, W. (2012). From Sensor Data to Animal Behaviour: An Oystercatcher Example. PLoS ONE, 7, e37997.

Shepard, E., Wilson, R., Halsey, L., Quintana, F., Gómez Laich, A., Gleiss, A., et al. (2008a). Derivation of body motion via appropriate smoothing of acceleration data. Aquatic Biology, 4, 235–241.

Shepard, E., Wilson, R., Quintana, F., Gómez Laich, A., Liebsch, N., Albareda, D., et al. (2008b). Identification of animal movement patterns using tri-axial accelerometry. Endangered Species Research, 10, 47–60.

Shuert, C.R., Pomeroy, P.P. & Twiss, S.D. (2018). Assessing the utility and limitations of accelerometers and machine learning approaches in classifying behaviour during lactation in a phocid seal. Animal Biotelemetry, 6.

Shuert, C.R., Pomeroy, P.P. & Twiss, S.D. (2020). Coping styles in capital breeders modulate behavioural trade-offs in time allocation: assessing fine-scale activity budgets in lactating grey seals (*Halichoerus grypus*) using accelerometry and heart rate variability. Behavioral Ecology and Sociobiology, 74.

Sönnichsen, L., Bokje, M., Marchal, J., Hofer, H., Jędrzejewska, B., Kramer-Schadt, S., et al. (2013). Behavioural Responses of European Roe Deer to Temporal Variation in Predation Risk. Ethology, 119, 233–243.

Stankowich, T. (2008a). Quantifying Behavior the JWatcher Way. Daniel T. Blumstein and Janice C. Daniel. Integrative and Comparative Biology, 48, 437–439.

Stankowich, T. (2008b). Ungulate flight responses to human disturbance: A review and meta-analysis. Biological Conservation, 141, 2159–2173.

Studd, E.K., Boudreau, M.R., Majchrzak, Y.N., Menzies, A.K., Peers, M.J.L., Seguin, J.L., et al. (2019a). Use of Acceleration and Acoustics to Classify Behavior, Generate Time Budgets, and Evaluate Responses to Moonlight in Free-Ranging Snowshoe Hares. Front. Ecol. Evol., 7.

Studd, E.K., Landry-Cuerrier, M., Menzies, A.K., Boutin, S., McAdam, A.G., Lane, J.E., et al. (2019b). Behavioral classification of low-frequency acceleration and temperature data from a free-ranging small mammal. Ecology and Evolution, 9, 619–630.

Takei, Y., Suzuki, I., Wong, M.K.S., Milne, R., Moss, S., Sato, K., et al. (2016). Development of an animal-borne blood sample collection device and its deployment for the determination of cardiovascular and stress hormones in phocid seals. American Journal of Physiology-Regulatory, Integrative and Comparative Physiology, 311, R788–R796.

Tatler, J., Cassey, P. & Prowse, T.A.A. (2018). High accuracy at low frequency: detailed behavioural classification from accelerometer data. The Journal of Experimental Biology, 221, jeb184085.

Tatler, J., Currie, S.E., Cassey, P., Scharf, A.K., Roshier, D.A. & Prowse, T.A.A. (2021). Accelerometer informed time-energy budgets reveal the importance of temperature to the activity of a wild, arid zone canid. Movement Ecology, 9.

Tixier, H. & Duncan, P. (1996). Are European roe deer browsers? A review of variations in the composition of their diets. Revue D’écologie.

Turner, D.C. (1979). An Analysis of Time-Budgeting by Roe Deer (*Capreolus capreolus*) in an Agricultural Area. Behaviour, 71, 246–290.

Tuyttens, F.A.M., de Graaf, S., Heerkens, J.L.T., Jacobs, L., Nalon, E., Ott, S., et al. (2014). Observer bias in animal behaviour research: can we believe what we score, if we score what we believe? Animal Behaviour, 90, 273–280.

Vehtari, A., Gelman, A. & Gabry, J. (2017). Practical Bayesian model evaluation using leave-one-out cross-validation and WAIC. Statistics and computing, 27, 1413–1432.

Volpov, B.L., Hoskins, A.J., Battaile, B.C., Viviant, M., Wheatley, K.E., Marshall, G., et al. (2015). Identification of Prey Captures in Australian Fur Seals (Arctocephalus pusillus doriferus) Using Head-Mounted Accelerometers: Field Validation with Animal-Borne Video Cameras. PLOS ONE, 10, e0128789.

Wang, Y., Nickel, B., Rutishauser, M., Bryce, C.M., Williams, T.M., Elkaim, G., et al. (2015). Movement, resting, and attack behaviors of wild pumas are revealed by tri-axial accelerometer measurements. Movement Ecology, 3.

Watanabe, S., Izawa, M., Kato, A., Ropert-Coudert, Y. & Naito, Y. (2005). A new technique for monitoring the detailed behaviour of terrestrial animals: A case study with the domestic cat. Applied Animal Behaviour Science, 94, 117–131.

Weimerskirch, H., Gault, A. & Cherel, Y. (2005). Prey distribution and patchiness: Factors in foraging success and efficiency of wandering albatrosses. Ecology, 86, 2611–2622.

Whitford, M. & Klimley, A.P. (2019). An overview of behavioral, physiological, and environmental sensors used in animal biotelemetry and biologging studies. Animal Biotelemetry, 7.

Whitney, N., Pratt, H., Pratt, T. & Carrier, J. (2010). Identifying shark mating behavior using three-dimensional acceleration loggers. Endangered Species Research, 10, 71–82.

Wijers, M., Trethowan, P., Markham, A., du Preez, B., Chamaillé-Jammes, S., Loveridge, A., et al. (2018). Listening to Lions: Animal-Borne Acoustic Sensors Improve Bio-logger Calibration and Behaviour Classification Performance. Front. Ecol. Evol., 6.

Wijers, M., Trethowan, P., du Preez, B., Chamaillé-Jammes, S., Loveridge, A.J., Macdonald, D.W., et al. (2021). The influence of spatial features and atmospheric conditions on African lion vocal behaviour. Animal Behaviour, 174, 63–76.

Wilmers, C.C., Nickel, B., Bryce, C.M., Smith, J.A., Wheat, R.E. & Yovovich, V. (2015). The golden age of bio-logging: how animal-borne sensors are advancing the frontiers of ecology. Ecology, 96, 1741–1753.

Wilson, R., Pütz, K., Grémillet, D., Culik, B. M., Kierspel, M., Regel, J., … & & Cooper, J (1995). Reliability of stomach temperature changes in determining feeding characteristics of seabirds. J Exp Biol, 198, 1115–1135.

Wilson, A.M., Lowe, J.C., Roskilly, K., Hudson, P.E., Golabek, K.A. & McNutt, J.W. (2013). Locomotion dynamics of hunting in wild cheetahs. Nature, 498, 185–189.

Wilson, R.P., Grundy, E., Massy, R., Soltis, J., Tysse, B., Holton, M., et al. (2014). Wild state secrets: ultra-sensitive measurement of micro-movement can reveal internal processes in animals. Frontiers in Ecology and the Environment, 12, 582–587.

Wilson, R.P., White, C.R., Quintana, F., Halsey, L.G., Liebsch, N., Martin, G.R., et al. (2006). Moving towards acceleration for estimates of activity-specific metabolic rate in free-living animals: the case of the cormorant: Activity-specific metabolic rate in free-living animals. Journal of Animal Ecology, 75, 1081–1090.

Wolf, M., van Doorn, G.S., Leimar, O. & Weissing, F.J. (2007). Life-history trade-offs favour the evolution of animal personalities. Nature, 447, 581–584.

Yoda, K., Sato, K., Niizuma, Y., Kurita, M., Bost, C., Le Maho, Y., et al. (1999). Precise monitoring of porpoising behaviour of Adélie penguins determined using acceleration data loggers. J Exp Biol, 202, 3121–3126.

