## Appendix for "Using accelerometers to infer behaviour of cryptic species in the wild"

**Table S1:** Ethogram of behaviours expressed by roe deer and used for this study.

| **5 Final behavioural classes** | **25 Behaviours regrouped for classification** | **54 Behaviours included in the initial ethogram** |
| --- | --- | --- |
| Running | Galloping | Galloping or leaping |
|  | Trotting | Trotting |
| Walking | Walking head-up | Walking head-up or head mid-height  Air sniffing or masticating head-up in motion |
| Foraging head-down | Walking head-down | Walking head-down  Ground sniffing or masticating head-down in motion |
|  | Feeding up head-down | Feeding up head-down |
|  | Unmoving up head down | Unmoving up head-down  Ground sniffing up or masticating head down  Drinking |
| Unmoving | Feeding head-up | Feeding head-up |
|  | Masticating head-up | Masticating or ruminating head-up |
|  | **Feeding down** | **Feeding down** |
|  | Masticating head down | Masticating or ***ruminating down*** |
|  | Unmoving up head up | Unmoving up head up |
|  | Observing up | Observing up, body or neck stretching up  Air smelling up |
|  | Vigilance up | Vigilance up |
|  | ***Unmoving down head down*** | ***Unmoving down head down or ground smelling down*** |
|  | Unmoving down head up | Unmoving down head up |
|  | Observing down | Observing or air smelling down  Neck stretching down |
|  | Vigilance down | Vigilance down |
|  | ***Urinating*** | ***Urinating*** |
|  | **Scratching the ground head down** | **Scratching the ground head down** |
|  | **Scratching the ground head up** | **Scratching the ground head up** |
|  | **Licking a congener** | **Licking a congener** |
|  | **Attacking** | **Threatening or butting** |
|  | **Vocalizing** | **Barking or squeaking** |
|  | ***Head smear*** | ***Head smear*** |
|  | **Getting up** | **Getting up** |
|  | ***Lie down*** | ***Lie down*** |
| Grooming | Grooming up | Grooming up |
|  | Scratching up head down | Scratching up head down |
|  | **Scratching up head up** | **Scratching up head up** |
|  | Snorting up | Snorting up |
|  | Head snorting up | Head snorting up |
|  | Grooming down | Grooming down |
|  | **Scratching down head down** | **Scratching down head down** |
|  | **Scratching down head up** | **Scratching down head up** |
|  | ***Head snorting down*** | ***Head snorting down*** |
| **Behaviours underrepresented or too brief (less than 2snd) in captive individuals *or only in captive individuals equipped with sensor B*** | | |

**Appendix S2: Behavioural sequences synchronised with acceleration data**

Initial inspection revealed that the internal clocks of the accelerometers often did not run at precisely the same rate as the GPS-beacon clock used during video-recording. This was particularly the case for the type A sensors, as the internal clock did not intermittently synchronize with the internal GPS system, causing a time drift in acceleration data (divergence = +/– 380 s for type A or +/– 6 s for type B). We thus corrected, when possible, the time of the accelerometers by aligning abrupt changes in the acceleration signal with abrupt changes in body movements of the observed animal (e.g. transition from walking to galloping, or observation to foraging head-down (Studd *et al*. 2019b)). Only acceleration data synchronised to behavioural sequences were used for further analyses (see Table S2).

**Table S2:** Details of the behavioural sequences per individual used in the Calibration and Validation steps

| **Procedure step** | **Individual** | **Sensor** | **Time of behavioural sequences included in the study** | **Behaviours absent in dataset among the 25 included in the study** | **Moment filmed** | **Moment used to scale acceleration data** |
| --- | --- | --- | --- | --- | --- | --- |
| Calibration | Fanny | A | 3h12 | Feeding head-up, Ruminting down, Head smear, Grooming down, Scratching up head-down, Unmoving down head-down, Vigilance down, Snorting head down | ≈ 1 week in January 2016 | Moment filmed on 2016-2017 |
|  | Ninon | A | 4h22 | Snorting head down, Head smear | ≈ 2 weeks in January-February 2017  ≈ 2 weeks in July 2017 |  |
|  | Ourale | A | 2h40 | Galloping, Snorting head up, Snorting, Total Unmoving up, Urinating, Lies down, Scratching up head-down, Vigilance down | ≈ 1 week in July 2017 |  |
|  | Fanny | B | 3h25 | Feeding head-up, Grooming down, Ruminating down, Urinating, Lies down, Unmoving down head-down, Vigilance down, Snorting head down, Head smear | ≈ 2 weeks in January 2019 | Moment filmed on 2019 |
|  | Ourale | B | 2h30 | Snorting head up, Scratching up head-down, Ruminating down, Urinating, Lies down, Unmoving down head-down, Vigilance down, Snorting head down, Head smear | ≈ 2 weeks in January 2019 |  |
| Validation | 980_17 | A | 20 min | Feeding head-up; Grooming down; snorting and snorting head down; head-smear; all unmoving behaviours down; lies down | 1 day in October (26/10)  3 days in November (8, 9 and 13/11) | Month of September  Month of September |
|  | 977_17 | A | 17 min | Feeding head-up; Grooming down, All snorting behaviours; head-smear; urinating; galloping and trotting, Scratching up head-down; Rumination down | 1 day in November (08/11) | Month of September |
|  | 978_17 | A | 25 min | Feeding head-up, All snorting behaviours; head-smear; urinating; trotting; Scratching up head-down; Unmoving down head-down; Total Unmoving up; Vigilance, Unmoving up head down | 1 day in October (26/10)  1 day in November (08/11) | Month of September  Month of September |
|  | 979_18 | A | 3 min | Feeding head-up; All grooming; All snorting behaviours; head-smear; all unmoving behaviours down; urinating; trotting; All unmoving behaviours up exepted vigilance; | 1 day in November (07/11) | Month of September |
|  | F1347_18 | A | 30 min | Feeding head-up; Grooming down, All snorting behaviours; head-smear; all unmoving behaviours down; | 2 days in October (30 and 31/10)  2 days in November (13 and 20/11) | Month of September  Month of September |
|  | 3023_18 | A | 26min30 | Grooming down, All snorting behaviours; head-smear; Scratching up head-down all unmoving behaviours down; Urinating; Lies down, trotting | 1 day in October (31/10)  3 days in November (07/11) | Month of September  Month of September |
|  | 3077_19 | B | 1h01 | Grooming down, All snorting behaviours; head-smear; all unmoving behaviours down; Scratching up head-down; Unmoving up head-down; urinating, Lies down, | 3 days in March (7,12 and 18/03)  1 day in May (01/05) | Month of March  Month of May |
|  | 3057_19 | B | 29 min | Grooming down, All snorting behaviours; head-smear; all unmoving behaviours down; urinating, Lies down. | 2 days in April (17 and 25/04)  1 day in May (21/05)  1 day in October (03/10) | Month of May  Month of May |
|  | 3075_19 | B | 51 min | Feeding head-up; Grooming down, snorting and snorting head down; head-smear; all unmoving behaviours down; Scratching up head-down; Unmoving up head-down; urinating, Lies down, | 1 day in March (12/03)  2 days in September (11 and 19/09)  1 day in november (15/11) | Month of March  Month of September  Month of September |
|  | 1005_19 | B | 12 min | All grooming behaviours; all snorting behaviours; head-smear; all unmoving behaviours down exepted Total unmoving down; Scratching up head-down; urinating; Lies down, | 1 day in April (03/04)  1 day in May (16/05) | Month of March  Month of May |
|  | 3073_19 | B | 28 min | Grooming down, snorting and snorting head down; head-smear; all unmoving behaviours down; Total unmoving up; urinating, Lies down, | 1 day in May (01/05)  2 days in September (11 and 12/09) | Month of May  Month of September |
|  | 3052_19 | B | 10min | All grooming behaviours; all snorting behaviours; head-smear; all unmoving behaviours down; Total unmoving up; urinating; Lies down, | 2 days in September (18 and 25/09) | Month of September |
|  | 887_19 | B | 1h44 | Grooming down, snorting head up and snorting head down; head-smear; all unmoving behaviours down; Total unmoving up; urinating, Lies down, | 3 days in April (17, 24 and 25/04))  4 days in May (8,9,14 and 22/05) | Month of May  Month of May |
|  | 3066_19 | B | 6 min | Feeding head-up; Foraging head-down; All Grooming behaviours, All snorting behaviours; head-smear; all unmoving behaviours down; Scratching up head-down; Total unmoving up; Ruminating up; Walking head-down; urinating, Lies down, | 2 days in March (07 and 12/03) | Month of March |

**Appendix S3: Metrics to evaluate algorithm performance**

For each algorithm, we compared the resulting predicted behaviours with the observed behaviours extracted from videos, based on confusion matrices. When considering a given behaviour against all others, these matrices provide true positives (TP, i.e. the number of acceleration values correctly assigned to a given behaviour), false positives (FP, i.e. the number of acceleration values incorrectly assigned to this behaviour), true negatives (TN, i.e. the number of acceleration values correctly assigned to a different behaviour) and false negatives (FN, i.e. the number of acceleration values incorrectly assigned to a different behaviour).

First, we evaluated algorithm performance for each behaviour separately, by calculating (1) precision, i.e. the proportion of observed acceleration values predicted as the true behaviour (2) recall (or sensitivity), i.e. the proportion of acceleration values for a predicted behaviour that were attributed to the true observed behaviour and (3) F1-score statistic, i.e. the harmonic mean of precision and recall. Secondly, we evaluated the overall performance of the algorithms by calculating by micro averaging (calculated on the overall confusion matrix (m)): (4) accuracy, i.e. the proportion of correct predictions and (5) the Kappa coefficient which compares the result of the classification versus a random classification. We also calculated by macro averaging (calculated on each behaviour and then averaged) the mean of (6) the Mathew correlation coefficient (MCC) (as an alternative to accuracy) and (7) the true skill Statistic (TSS) (as an alternative to Kappa) which are less sensitive to sample size (Pagan*o et a*l. 2017; Tatle*r et a*l. 2018). As the MCC metric is undefined when a whole row or column of the confusion matrix is zero, we replaced any zeros by 1 (Chicco & Jurman 2020). We compared values of MCC obtained by this correction method with those obtained by the method proposed by Chicc*o et a*l. 2021, inferring a MCC value of -1, 0 or 1 according to the value of the confusion matrix (TP, TN, FP or FN) which is zero (see Table S3).

Metrics (1:5) range between [0, 1], while metrics (6) and (7) range between [−1, +1].

1) Precision =
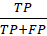

2) Sensitivity =
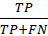

3) F1-score = 
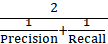

4) Accuracy =
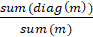

5) Kappa=
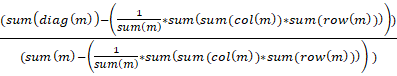

6) MCC = ∑
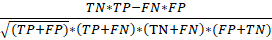

7) TSS =∑
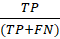
+
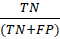
-1

**Table S3:** Comparison of global performances of different machine learning algorithms on the captive dataset (Calibration Step), estimated by accuracy, kappa, Mean (MCC) and Mean (TSS) expressed in percentage (%), for the 25 behaviours and the 5 behavioural classes. We added Mean (MCC) values calculated following Chicco *et al*. (2021) in grey for comparison with those obtained with Chicco & Jurman (2020)’s method.

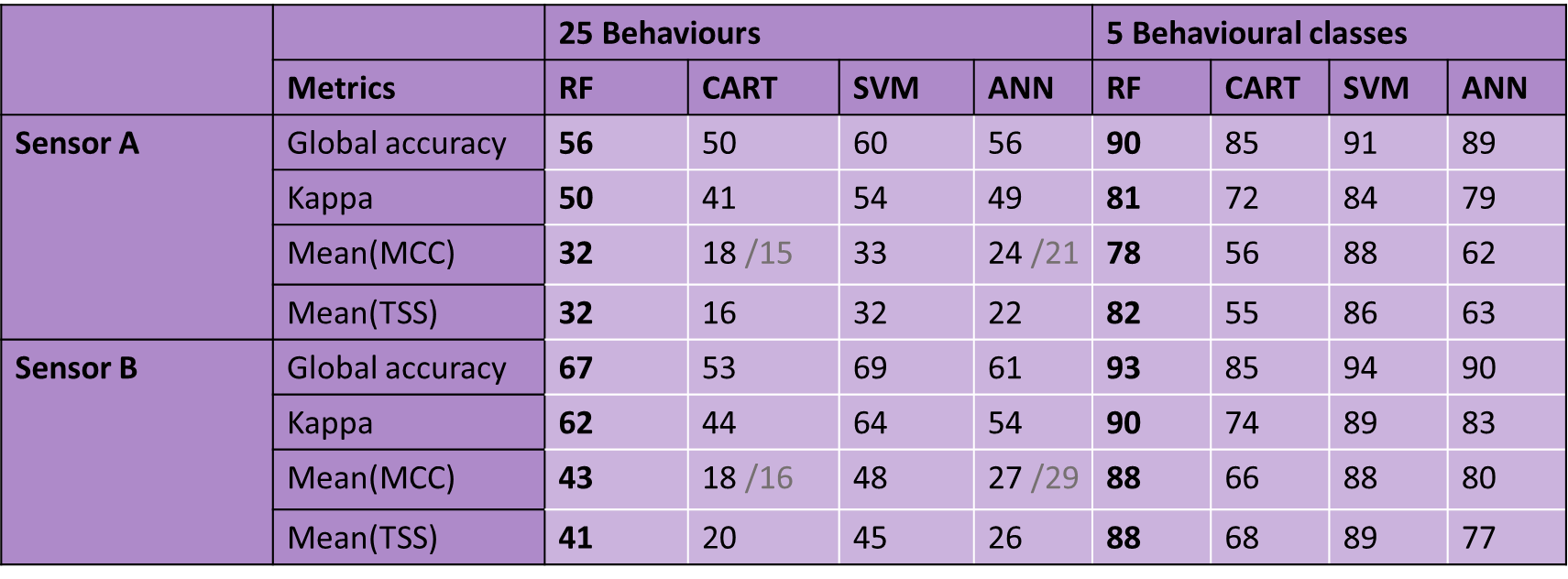

**Table S4.a:** Confusion matrix between behaviours predicted by the Random Forest algorithm (rows) and observed behaviours (columns) in captive roe deer equipped with sensor A (Calibration Step). Performance metrics (Sensitivity, Precision and F1-score expressed in percentage (%)) are also provided. Correct classifications are in **bold**.

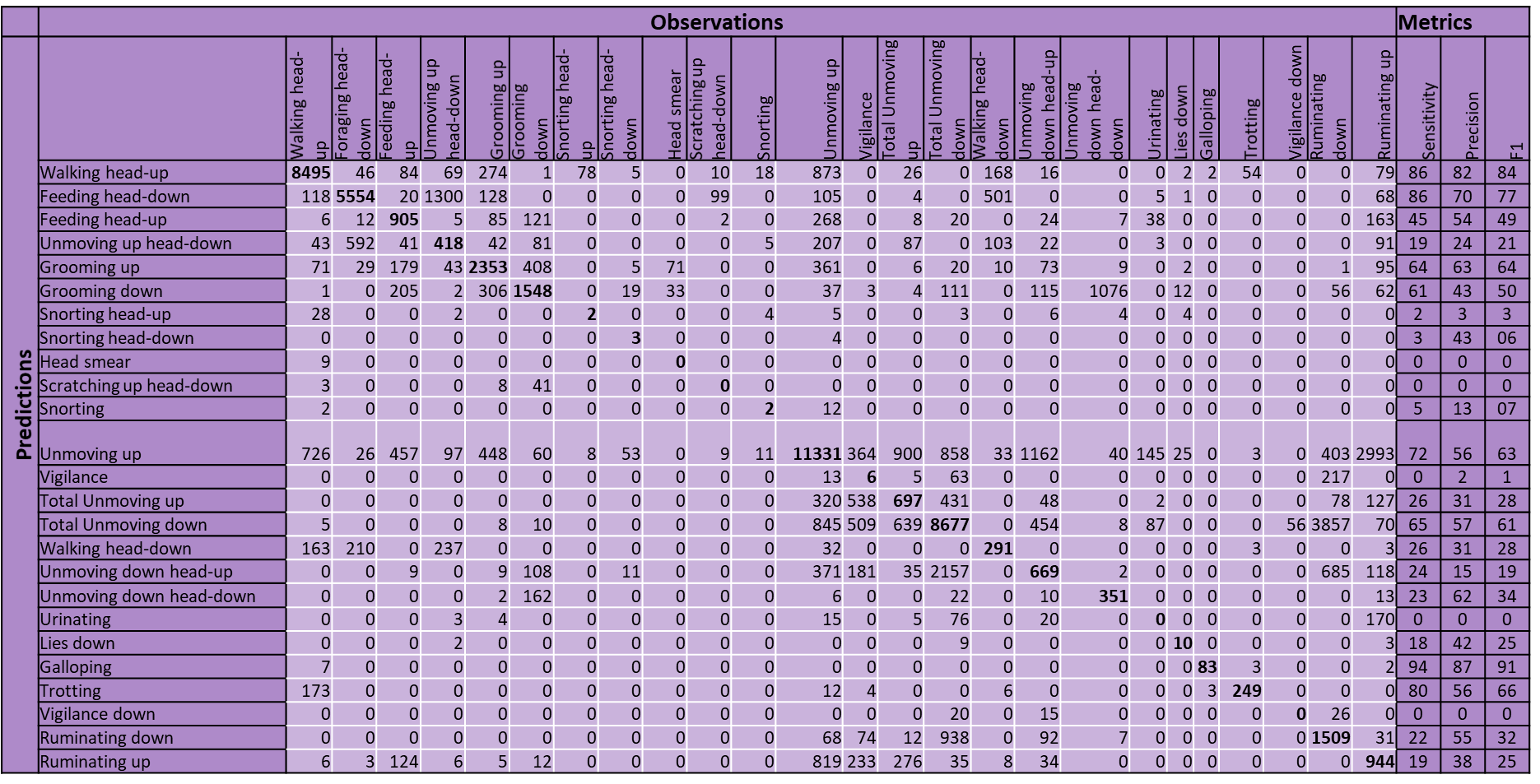

**Table S4.b:** Confusion matrix between behaviours predicted by the Random Forest algorithm (rows) and observed behaviours (columns) in captive roe deer equipped with sensor B (Calibration Step). Performance metrics (Sensitivity, Precision and F1-score, expressed in percentage (%)) are also provided. Correct classifications are in **bold**.

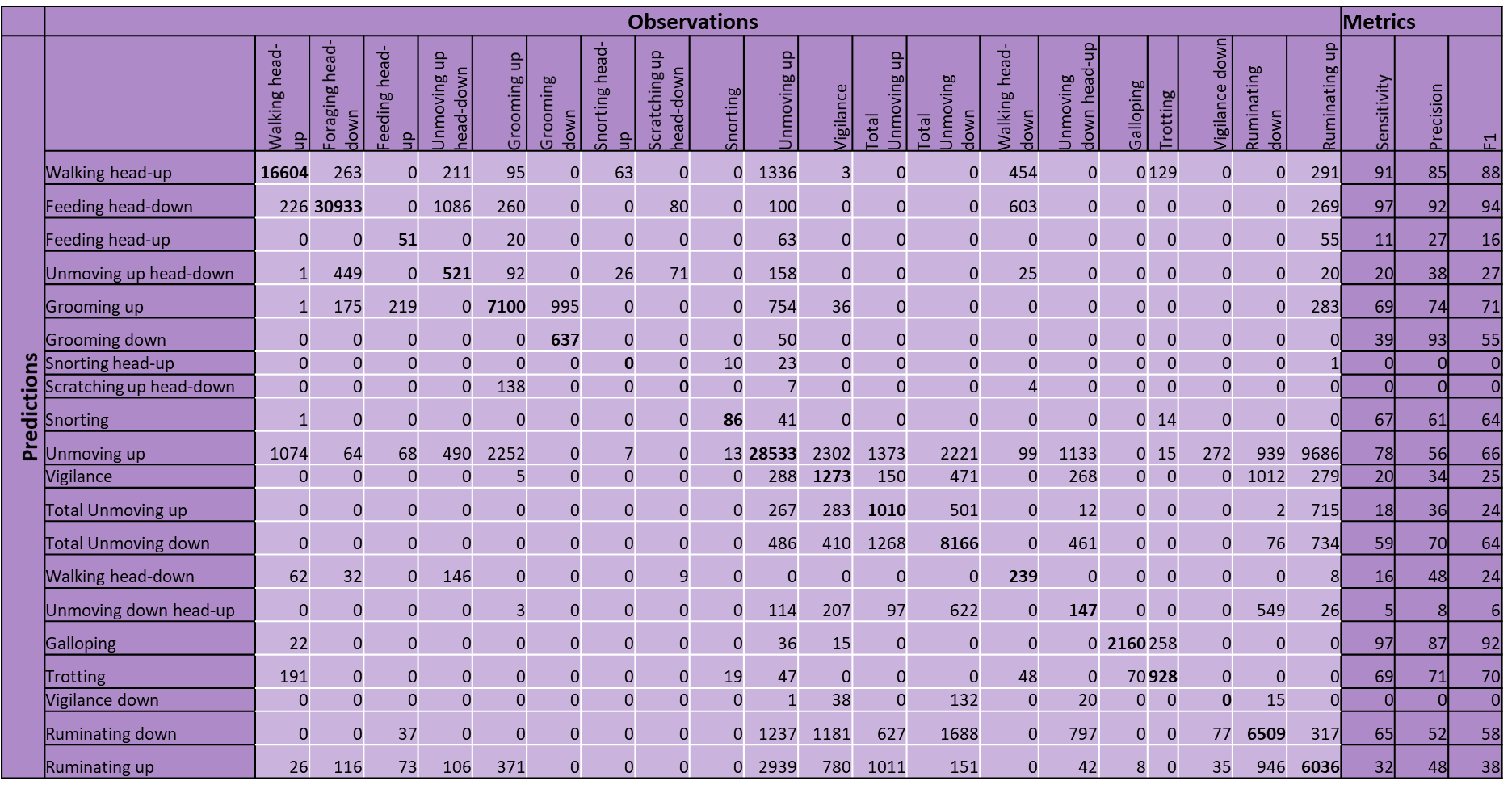

**Table S5:** Comparison of performance metrics for the 5 behavioural classes assessed by several machine learning methods applied on captive roe deer (Calibration Step). The performance of each behavioural class was estimated by sensitivity, precision and F1-score, expressed in percentage (%).

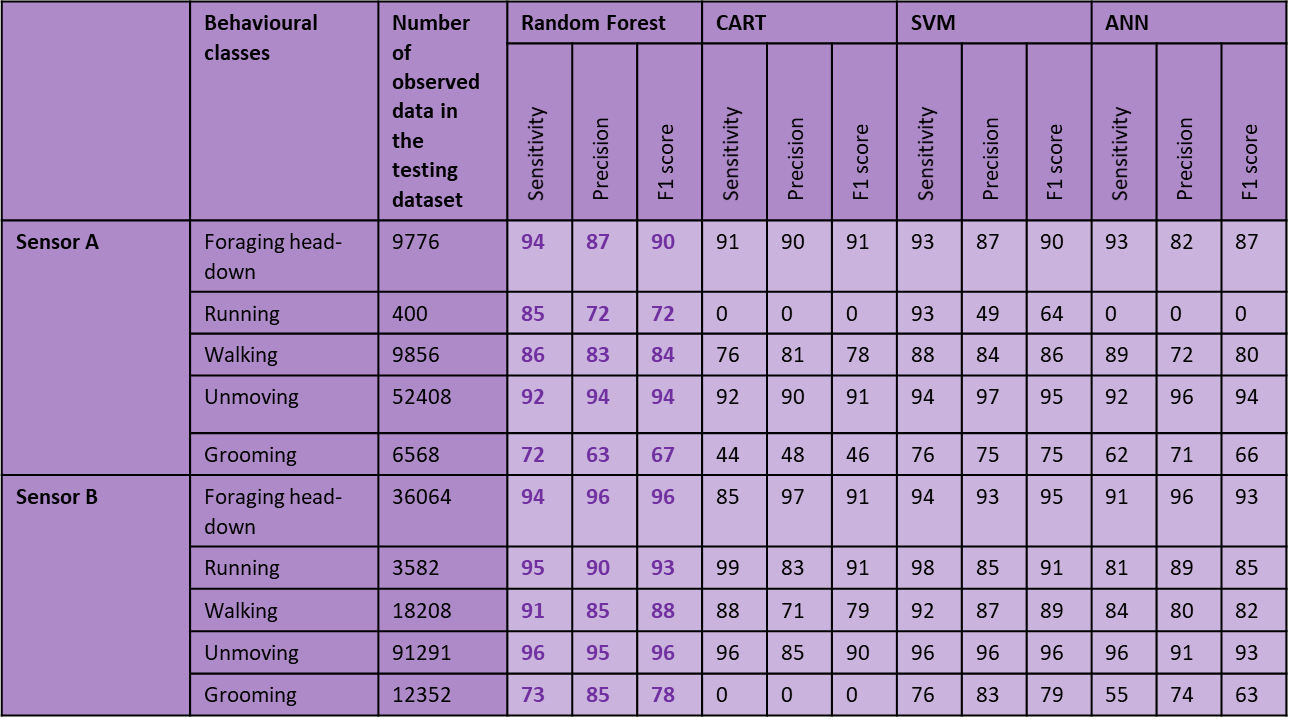

**Appendix S6: Accelerometer variables to infer behaviours using random forest algorithms**

Variable importance varied markedly between sensor types (A or B) (e.g. see Table S6 for global algorithms), presumably because of the different orientations of their accelerometric axes. These accelerometer-related variables also allowed us to clearly discriminate the five behavioural classes for both sensors. For instance, for type A sensors, high mean values of pitch and static[Y] were indicative of a head-down posture, mostly corresponding to the foraging head-down behaviour. Walking and, particularly, running behaviours had high mean values of SD[X] and SD[Y] and mDBA[X], which were indicative of forward-backward motion. Finally, grooming had a higher amplitude for the values of yaw, indicative of head movements with a large amplitude to the right or left (see Table S7.a). Similarly, for type B sensors, foraging head-down had high mean values of roll and static[X]. Walking and, particularly, running had high mean values of SD[Y] and mDBA[Y], but low mean values of minDBA[Y], whereas grooming had a higher amplitude for values of static[Z] (Table S7.b).

Although values of OBBA and VEDBA were not particularly informative variables for behavioural discrimination, they are often used to index energy expenditure (Qase*m et a*l. 2012). Thus, it is interesting to note they varied consistently among behavioural classes with high values for running, then walking, and lower values for foraging head-down, grooming and, lastly, unmoving (Tables S7).

**Figure S6:** Variable importance in the global Random Forest algorithms: a) Sensor A; b) Sensor B. The importance plot provides a relative ranking of parameters in which higher values indicate parameters that contributed more toward classification performance.

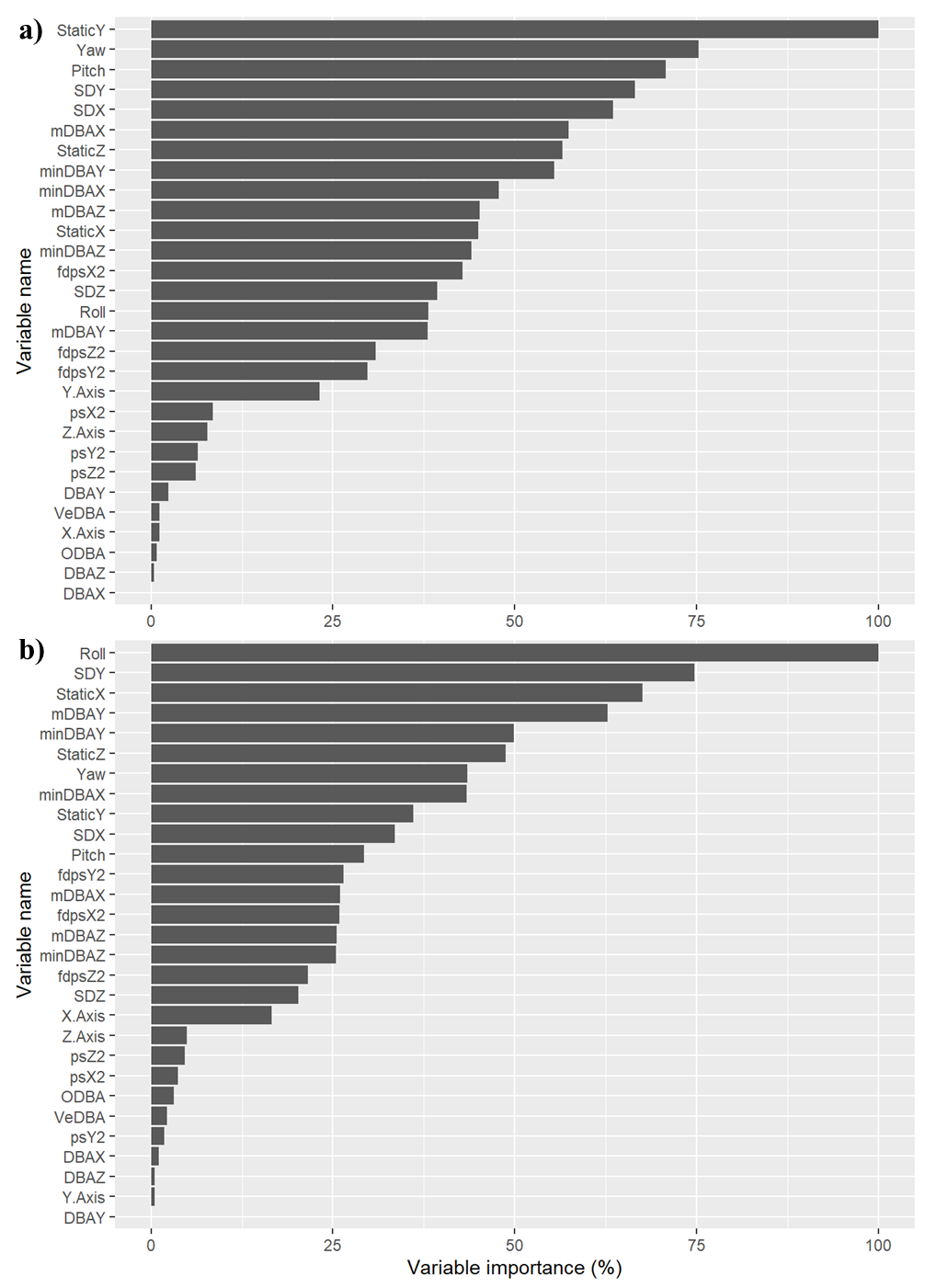

**Table S7:** Description of value changes for different variables of Random Forest algorithms, (a) for sensor A and (b) for sensor B, used to discriminate the behavioural classes.

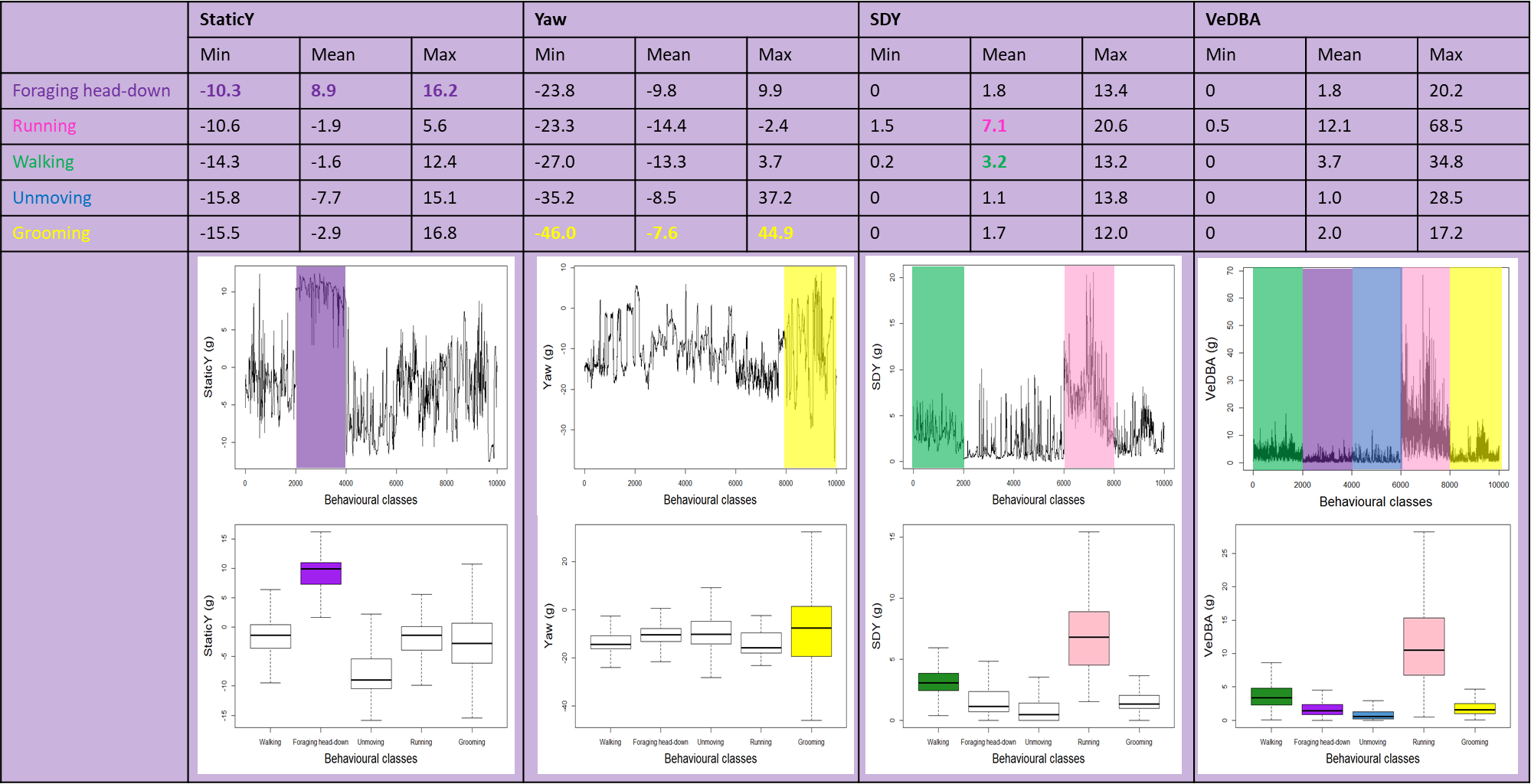

A)

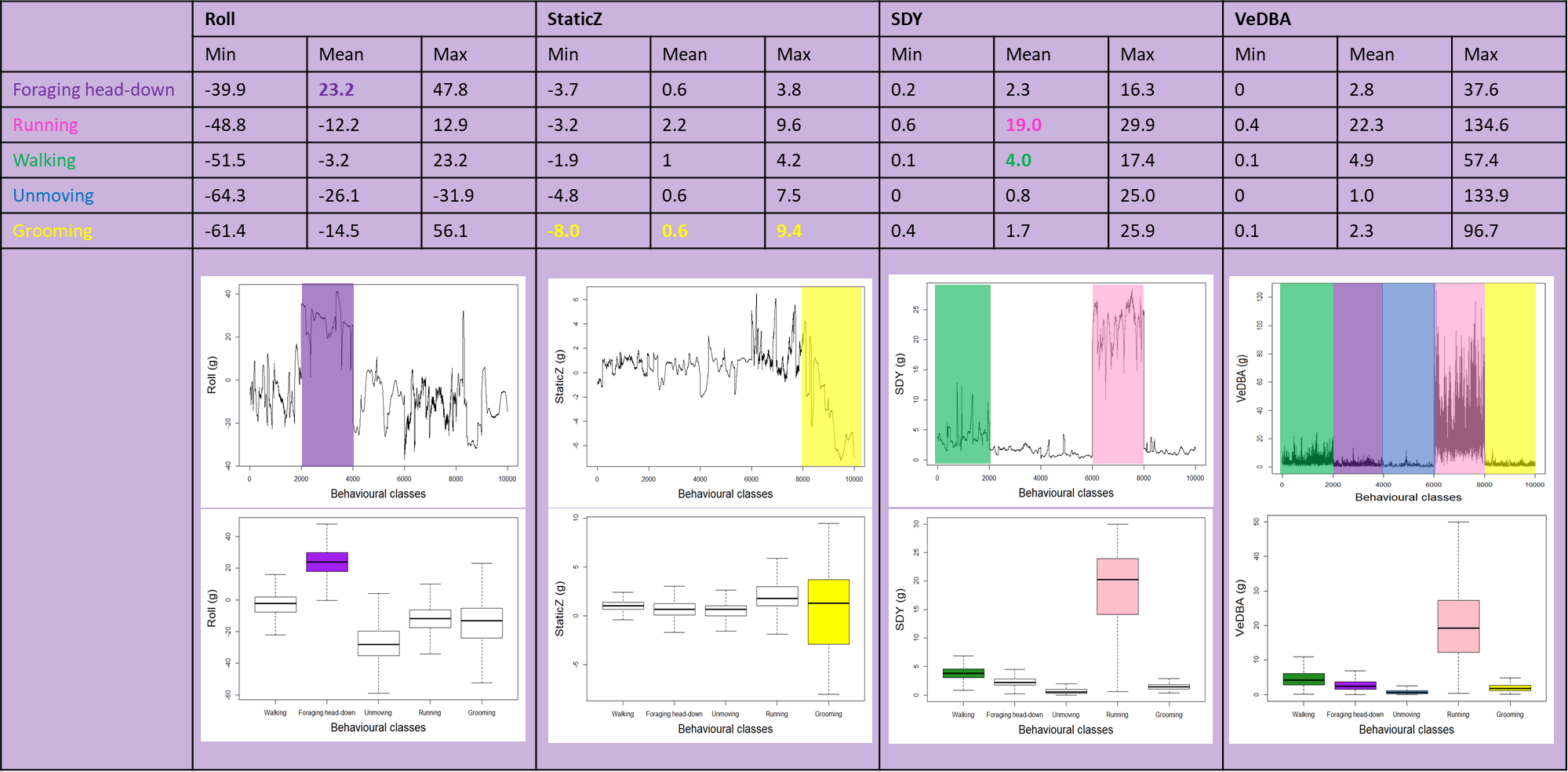

Table 3 : Variables derived from accelerometer data used to the classification of behaviours

B)

**Table S8**: Comparison of global performances of different machine learning algorithms on the free-ranging dataset (Validation Step), estimated by accuracy, kappa, Mean (MCC) and Mean (TSS) expressed in percentage (%), for the 25 behaviours and the 5 behavioural classes. We added Mean (MCC) values calculated following Chicco *et al*. (2021) in grey for comparison with those obtained with Chicco & Jurman (2020)’s method.

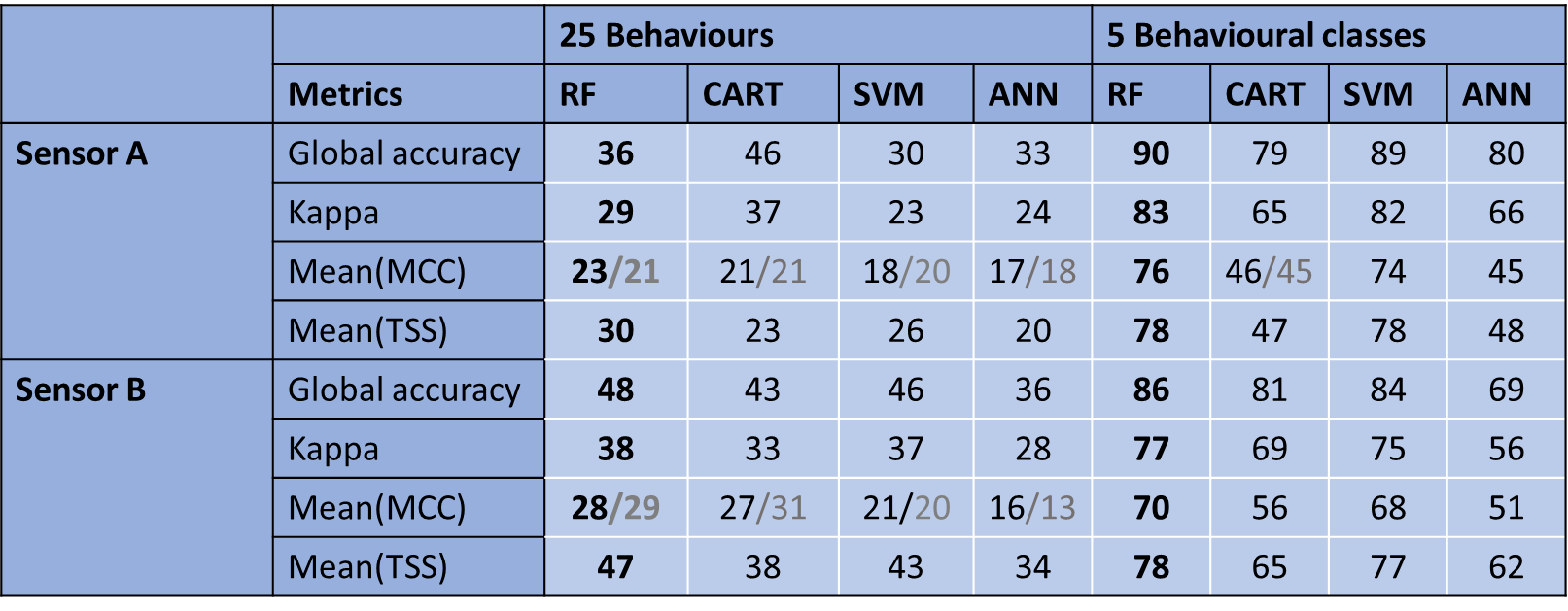

**Table S9.a:** Confusion matrix between behaviours predicted by the Random Forest algorithm (rows) and observed behaviours (columns) in free-ranging roe deer equipped with sensor A (Validation Step). Performance metrics (Sensitivity, Precision and F1-score expressed in percentage (%)) are also provided. Correct classifications are in **bold**.

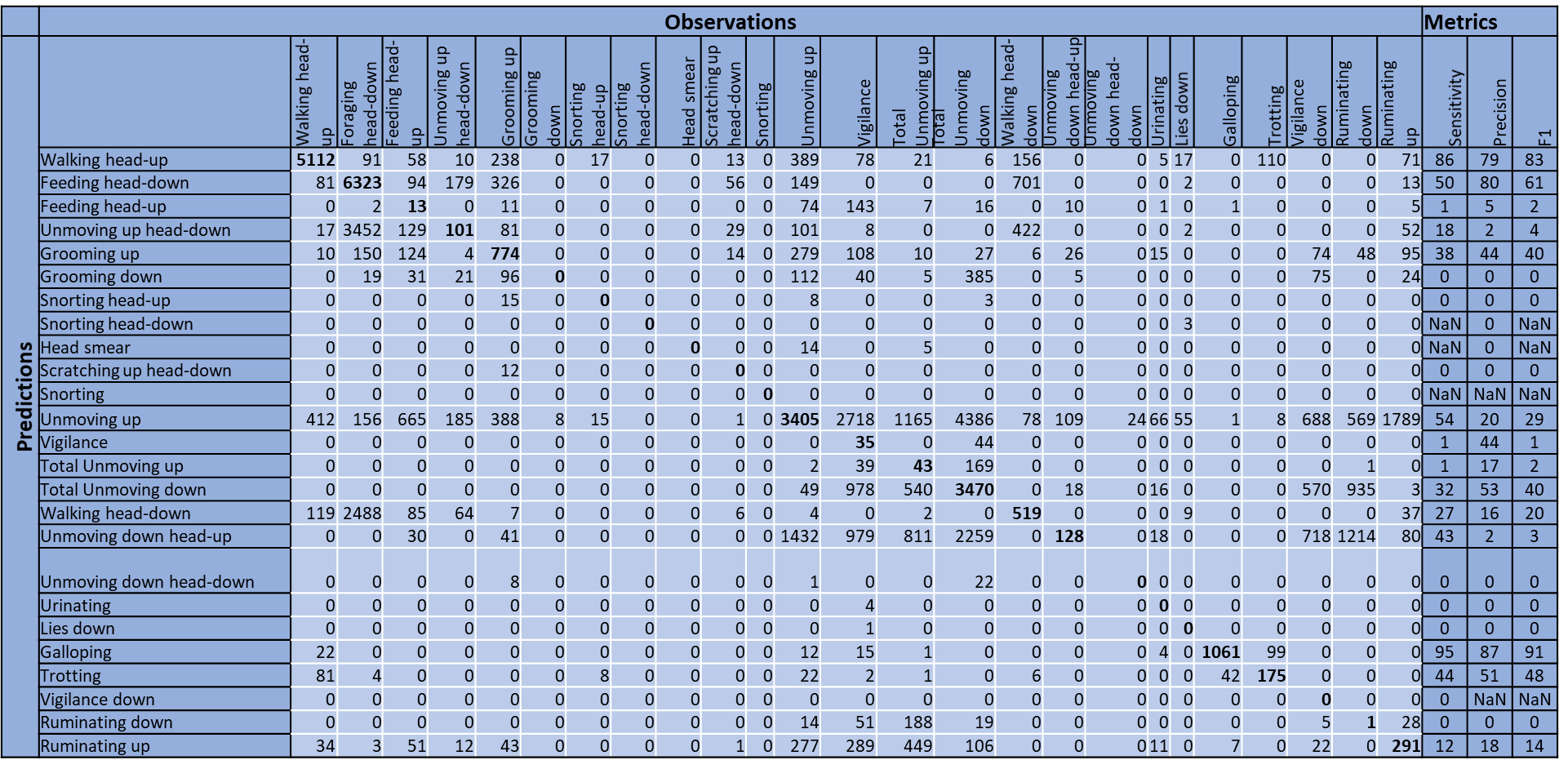

**Table S9.b:** Confusion matrix between behaviours predicted by the Random Forest algorithm (rows) and observed behaviours (columns) in free-ranging roe deer equipped with sensor B (Validation Step). Performance metrics (Sensitivity, Precision and F1-score, expressed in percentage (%)) are also provided. Correct classifications are in **bold**.

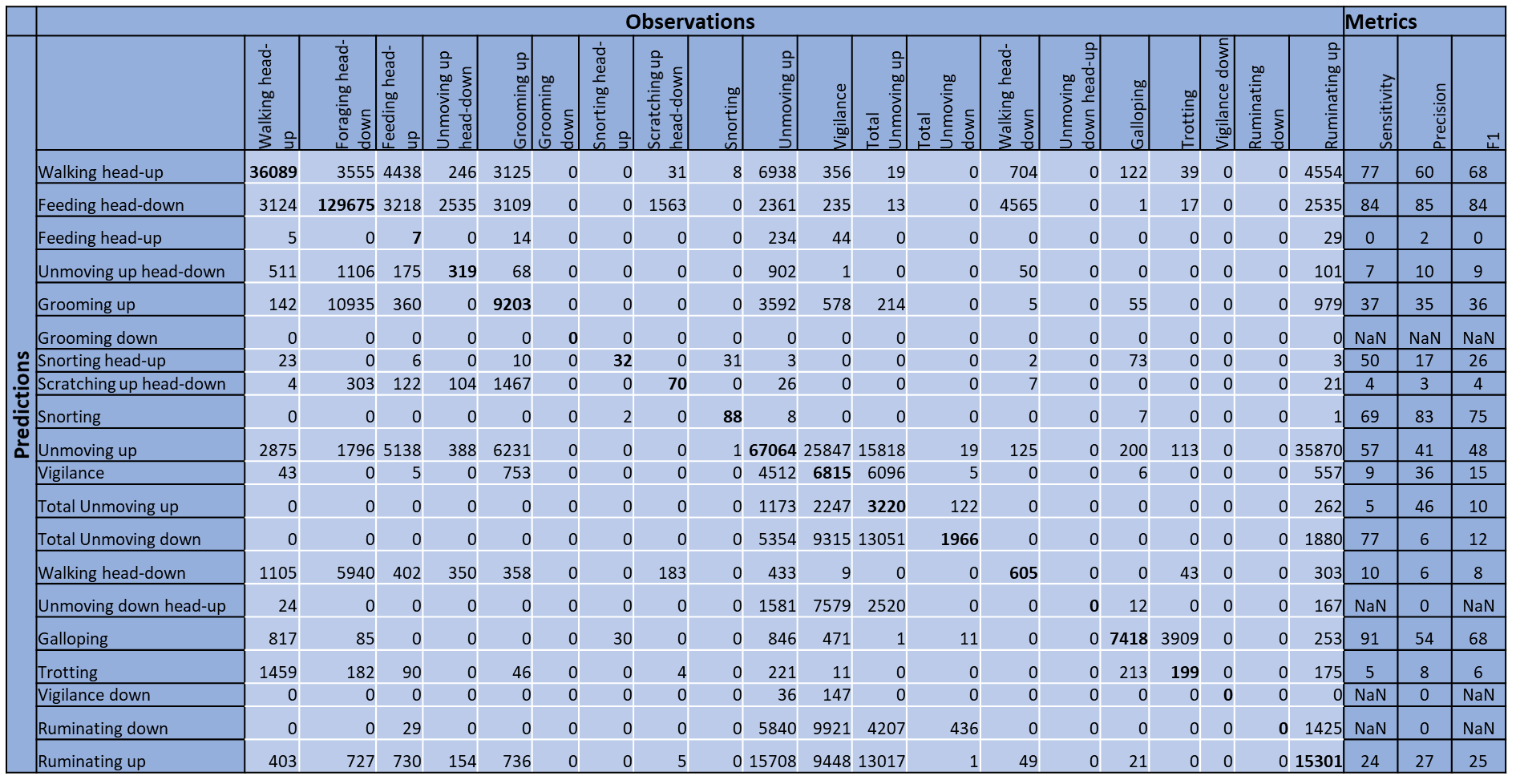

**Table S10:** Comparison of performance metrics for the 5 behavioural classes assessed by several machine learning methods applied on free-ranging roe deer (Validation Step). The performance of each behavioural class was estimated by sensitivity, precision and F1-score, expressed in percentage (%).

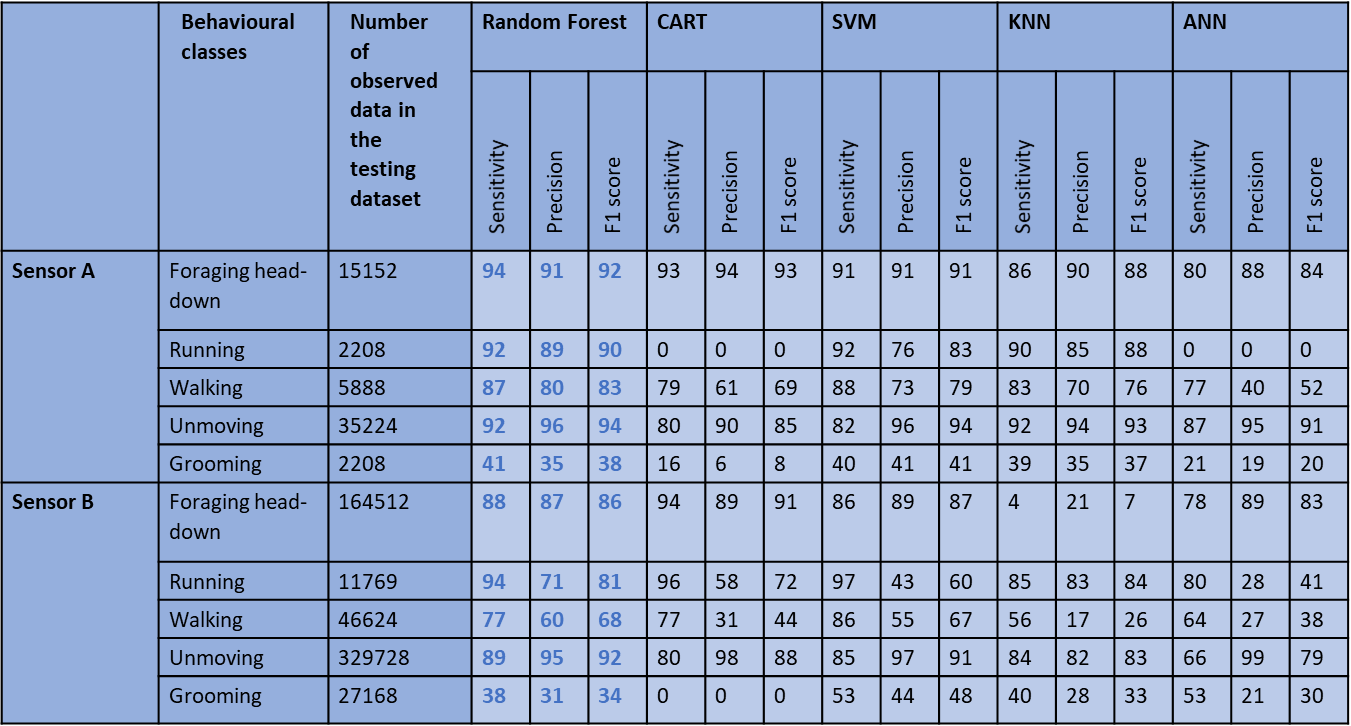

**Figure S11:** Example of comparison of observed (top) and predicted (down) behavioural classes along a) X.axis, b) Y.Axis and c) Z.Axis in the validation step, for a free-ranging individual that was video-recorded.

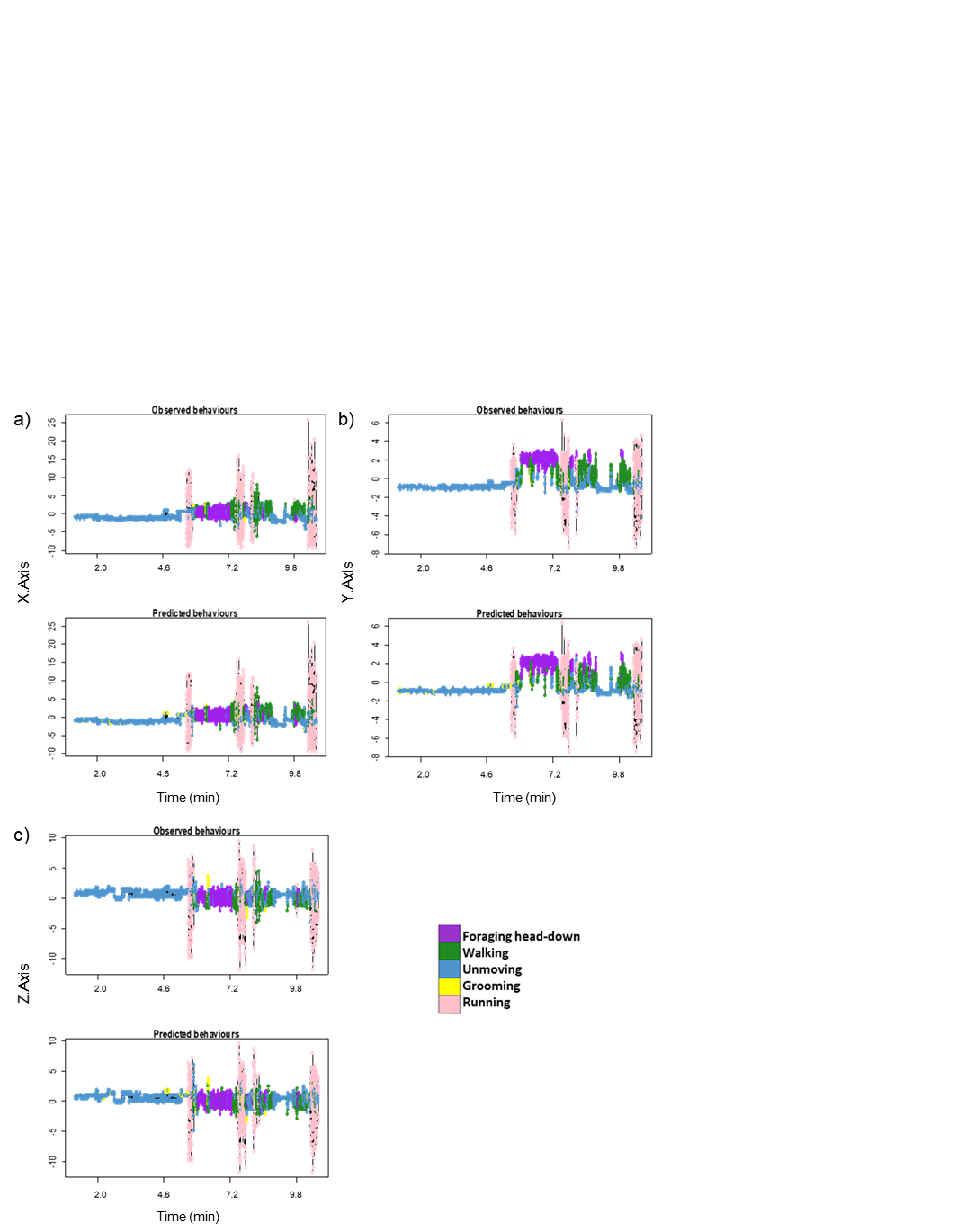
